# RUNX1 safeguards the identity of the fetal ovary through an interplay with FOXL2

**DOI:** 10.1101/598607

**Authors:** Barbara Nicol, Sara A. Grimm, Frederic Chalmel, Estelle Lecluze, Maëlle Pannetier, Eric Pailhoux, Elodie Dupin-De-Beyssat, Yann Guiguen, Blanche Capel, Humphrey H.-C. Yao

## Abstract

Sex determination of the gonads begins with fate specification of gonadal supporting cells into either ovarian granulosa cells or testicular Sertoli cells. This process of fate specification hinges on a balance of transcriptional control. We discovered that expression of the transcription factor RUNX1 is enriched in the fetal ovary in rainbow trout, turtle, mouse, goat and human. In the mouse, RUNX1 marks the supporting cell lineage and becomes granulosa cell-specific as the gonads differentiate. RUNX1 plays complementary/redundant roles with FOXL2 to maintain fetal granulosa cell identity, and combined loss of RUNX1 and FOXL2 results in masculinization of the fetal ovaries. At the chromatin level, RUNX1 occupancy overlaps partially with FOXL2 occupancy in the fetal ovary, suggesting that RUNX1 and FOXL2 target a common set of genes. These findings identify RUNX1, with an ovary-biased pattern conserved across species, as a novel regulator in securing the identity of ovarian supporting cells and the ovary.

---

A critical step that shapes the reproductive identity of the embryo is the sexual differentiation of the bipotential gonads. Supporting cells in the fetal gonads are the first cell population to differentiate, and dictate the fate of the gonads. As a consequence, defects in supporting cell differentiation have dire consequences on reproductive outcomes of the individual, from sex-reversal to infertility. Supporting cells differentiate into either Sertoli cells, which drive testis development, or granulosa cells, which control ovarian development. It has become clear that supporting cell differentiation, and maintenance of their commitment, requires a coordinated action of multiple factors that play either complementary, redundant and even antagonistic roles^1^. For instance, fate decision and maintenance of ovarian identity relies mainly on two conserved elements: the WNT4/RSPO1/beta-catenin pathway^2, 3, 4^ and the transcription factor FOXL2^5, 6, 7^. These two elements synergistically promote expression of pro-ovarian genes and at the same time, antagonize key pro-testis factors such as SOX9 and DMRT1. However, the combined loss of these two key pro-ovarian signaling only results in an incomplete inactivation of ovarian differentiation, suggesting that additional pro-ovarian factors are at play during gonadal differentiation^8, 9^. Factors involved in gonad differentiation are generally conserved in vertebrates and even invertebrates, although their position in the hierarchy of the molecular cascade may change during evolution^10^. For instance, the pro-ovarian transcription factor FOXL2 is important for ovarian differentiation/function in human^11^, goat^12^ and fish^13, 14^. The pro-testis transcription factor DMRT1 is highly conserved and critical for testis development in worms, fly^15^, fish^16, 17^ and mammals^18, 19, 20^.

In this study, we set up to investigate the role of transcription factor RUNX1 in the mouse fetal ovary. In *Drosophila melanogaster*, the *RUNX* ortholog *runt* is essential for ovarian determination^21, 22^. In the mouse, *Runx1* mRNA is enriched in the fetal ovary based on transcriptomic analyses^23^. The RUNX family arose early in evolution^24^: members have been identified in metazoans from sponge to human, where they play conserved key roles in developmental processes. In vertebrates, RUNX1 acts as a transcription factor critical for cell lineage specification in multiple organs, and particularly in cell populations of epithelial origin^25^. We first established the expression profile of *RUNX1* in the fetal gonads in multiple vertebrate species from fish to human. We then used knockout mouse models and genomic approaches to determine the function and molecular action of RUNX1 and its interplay with another conserved ovarian regulator, FOXL2, during supporting cell differentiation in the fetal ovary.

## Results

### The pattern of *Runx1* expression implies a potential role in fetal ovarian differentiation

The *runt* gene, critical for ovarian differentiation in the fly^21^, has 3 orthologs in mammals: *RUNX1*, *RUNX2* and *RUNX3*. While all three RUNX transcription factors bind the same DNA motif, they are known to have distinct, tissue-specific functions^26^. In the mouse, *Runx1* was the only one with a strong expression in the fetal ovary whereas *Runx2* and *Runx3* were expressed weakly in the fetal gonads in a non-sexually dimorphic way (Fig. 1a). At the onset of sex determination (Embryonic day or E11.5), *Runx1* expression was similar in both fetal testis and ovary before becoming ovary-specific after E12.5 (Fig. 1b), consistent with observations by others^23^. An ovary-enriched expression of *Runx1* during the window of early gonad differentiation was also observed in other mammals such as human and goat, as well as in species belonging to other classes of vertebrates such as the red-eared slider turtle and rainbow trout (Fig. 1c-f), implying an evolutionarily conserved role of RUNX1 in ovary differentiation.

**Figure 1:**
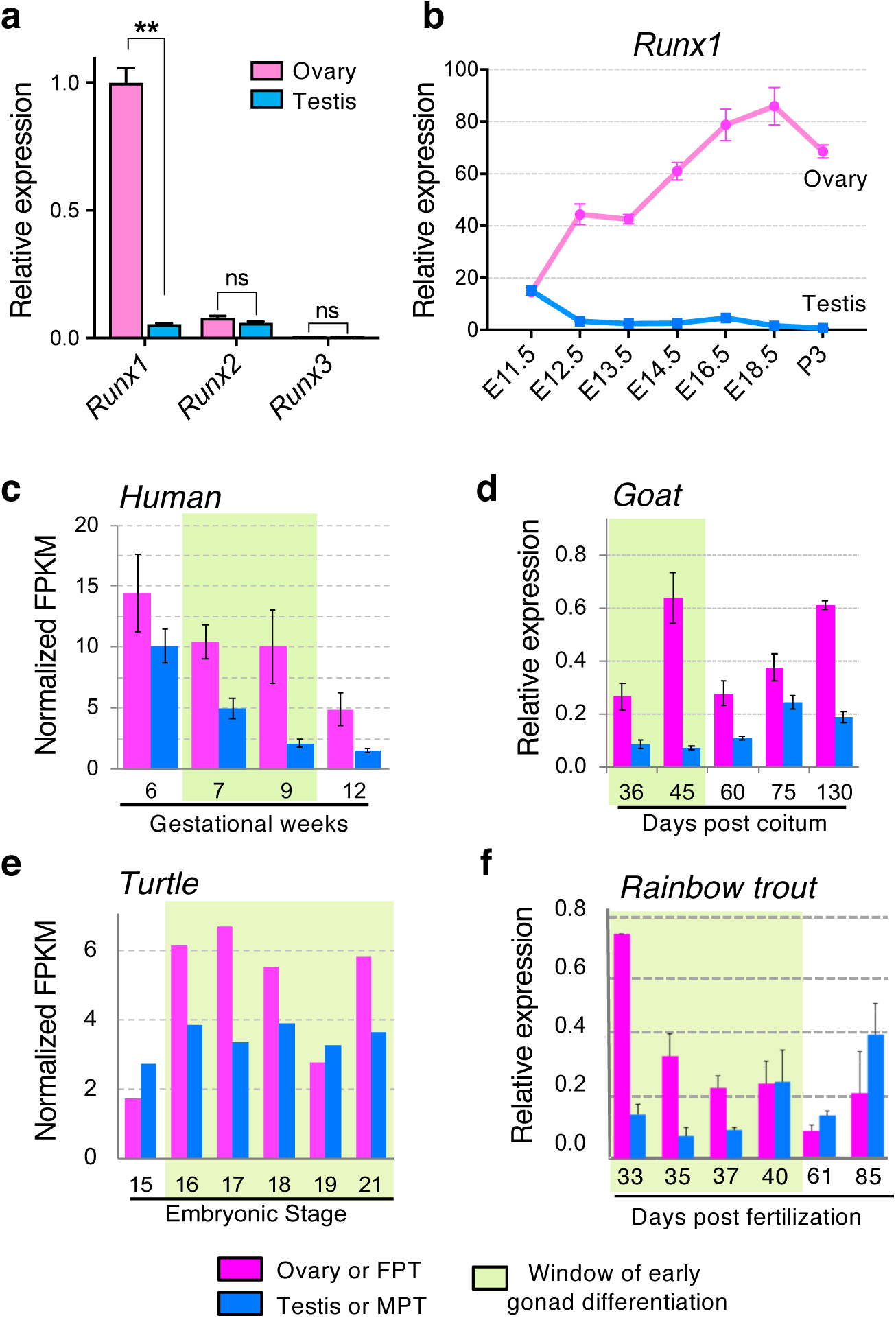
*RUNX1* expression is enriched in female gonads during gonadal differentiation in the mouse and other vertebrates. **(a)** Expression of *Runx1*, *Runx2* and *Runx3* mRNAs in ovaries and testis of E14.5 mouse embryos (n= 5/sex); Values are presented as mean ± s.e.m.; non-parametric t-test, \*\**P*<0.01; ns: not significant. **(b)** Expression time course of *Runx1* mRNA in mouse gonads during gonadal differentiation (n=3/stage); Values are presented as mean ± s.e.m. **(c-f)** Time course of *RUNX1* mRNA expression in four other vertebrate species, human, goat, red-eared slider turtle and rainbow trout during gonad differentiation. For the turtle, pink and blue bars represent gonads at Female-Promoting Temperature FPT of 31°C and at Male-Promoting Temperature MPT of 26°C respectively^62^. *RUNX1* expression was analyzed by RNA-seq in human and red-eared slider turtle^62^, and by qPCR in the goat and rainbow trout. Green highlighted areas represent the window of early gonadal differentiation.

To identify the cell types that express *Runx1* in the gonads, we examined a reporter mouse model that produces EGFP under the control of *Runx1* promoter^27^ (Fig. 2). Consistent with the time-course of *Runx1* mRNA expression (Fig. 1b), *Runx1*-EGFP was present in both fetal ovary and testis at E11.5, and then increased in the ovary and diminished in the testis at E12.5 onwards (Fig. 2). At E11.5 in both testis and ovary, *Runx1*-EGFP was present in a subset of SF1+/PECAM-somatic cell population whereas absent in the SF1-/PECAM+ germ cells (Fig. 2a-d). In the testis, these *Runx1*- EGFP+ somatic cells corresponded to Sertoli cells, as demonstrated by a complete overlap with SRY, the sex-determining factor that drives Sertoli cell differentiation^28^ (Fig. 2e and S1). At this stage, there is no marker for ovarian supporting cells that allow us to determine which subset of somatic cells were positive for *Runx1*-EGFP in the ovary. However, at E12.5, when the sex of gonads becomes morphologically distinguishable, *Runx1*-EGFP was specifically detected in the supporting cell lineage of both sexes: strongly in FOXL2+ pre-granulosa cells of the ovary (Fig. 2g), and weakly in SOX9+ Sertoli cells of the testis (Fig. 2f and S1). *Runx1*-EGFP expression was eventually turned off in the fetal testis while maintained in the ovary (Fig. 2h & i). Throughout fetal development of the ovary, *Runx1*-EGFP remained in FOXL2+ pre-granulosa cells (Fig. 3). *Runx1*-EGFP was also detected in the ovarian surface epithelium at E16.5 and birth (arrows in Fig. 3b-c), which gives rise to granulosa cells in the cortex of the ovary ^29, 30^. *Runx1*-EGFP was also expressed in somatic cells of the cortical region right underneath the surface epithelium, and some of these *Runx1*-EGFP+ cells presented a weak expression of FOXL2 (Fig. 3g-i, arrowheads). In summary, *Runx1* marks the supporting cell lineage in the gonads at the onset of sex determination, and becomes granulosa cell-specific as gonads differentiate.

**Figure 2:**
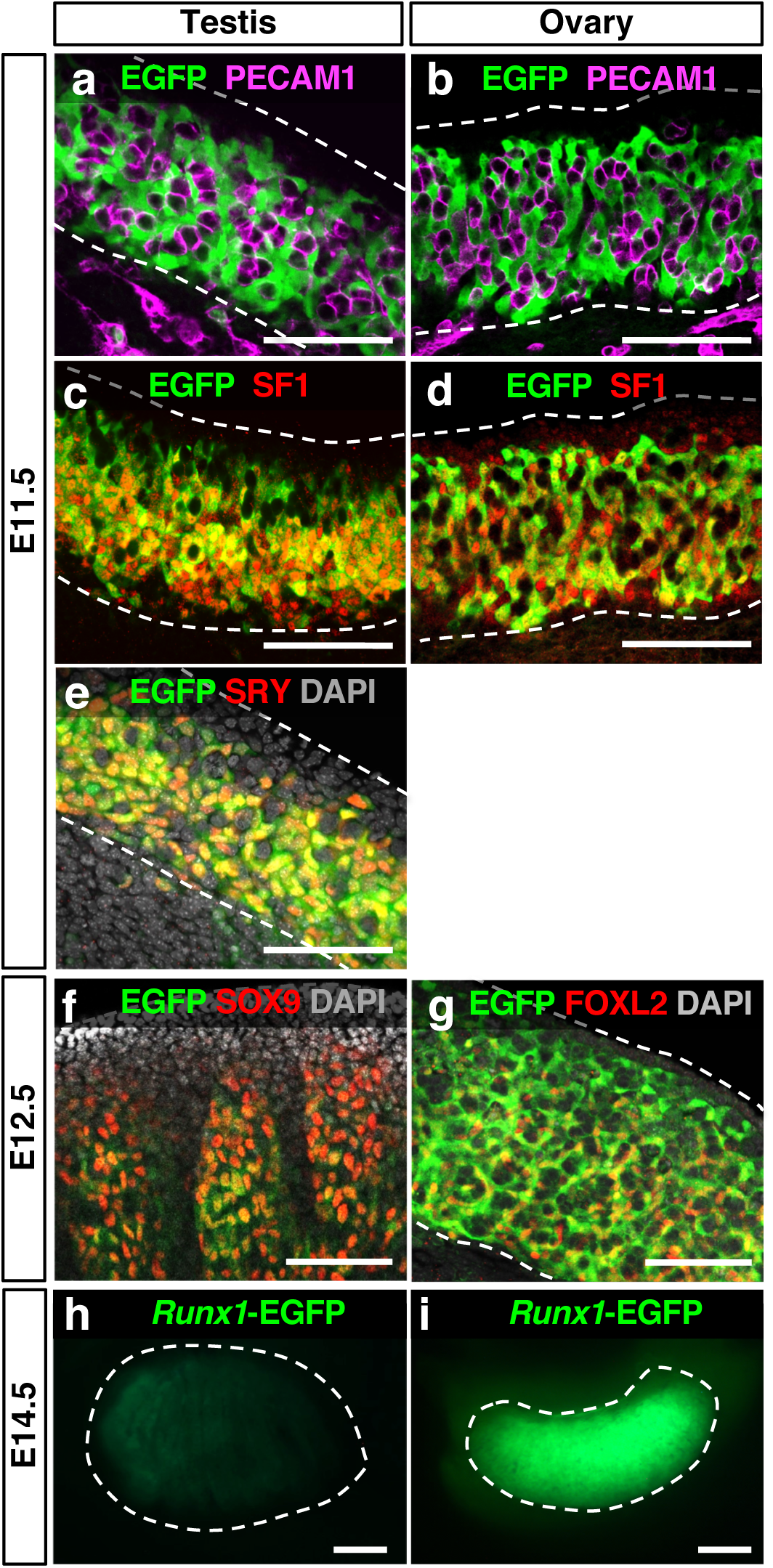
*Runx1* is expressed in the supporting cell lineage during gonad differentiation in the mouse embryos. **(a-g)** Whole mount immunofluorescence of testes and ovaries from Tg(Runx1-EGFP) reporter mice at E11.5 and E12.5. Gonads with endogenous EGFP were co-labeled with markers for germ cells/vasculature (PECAM-1; a-b), somatic cells (SF1; c-d), Sertoli cells in the testis (SRY in e and SOX9 in f), and for granulosa cells in the ovary (FOXL2; g). Scale bars: 100 μm. **(h-i)** Detection of endogenous EGFP in freshly collected E14.5 gonads. Scale bars: 200 μm. Dotted lines outline the gonads.

**Figure 3:**
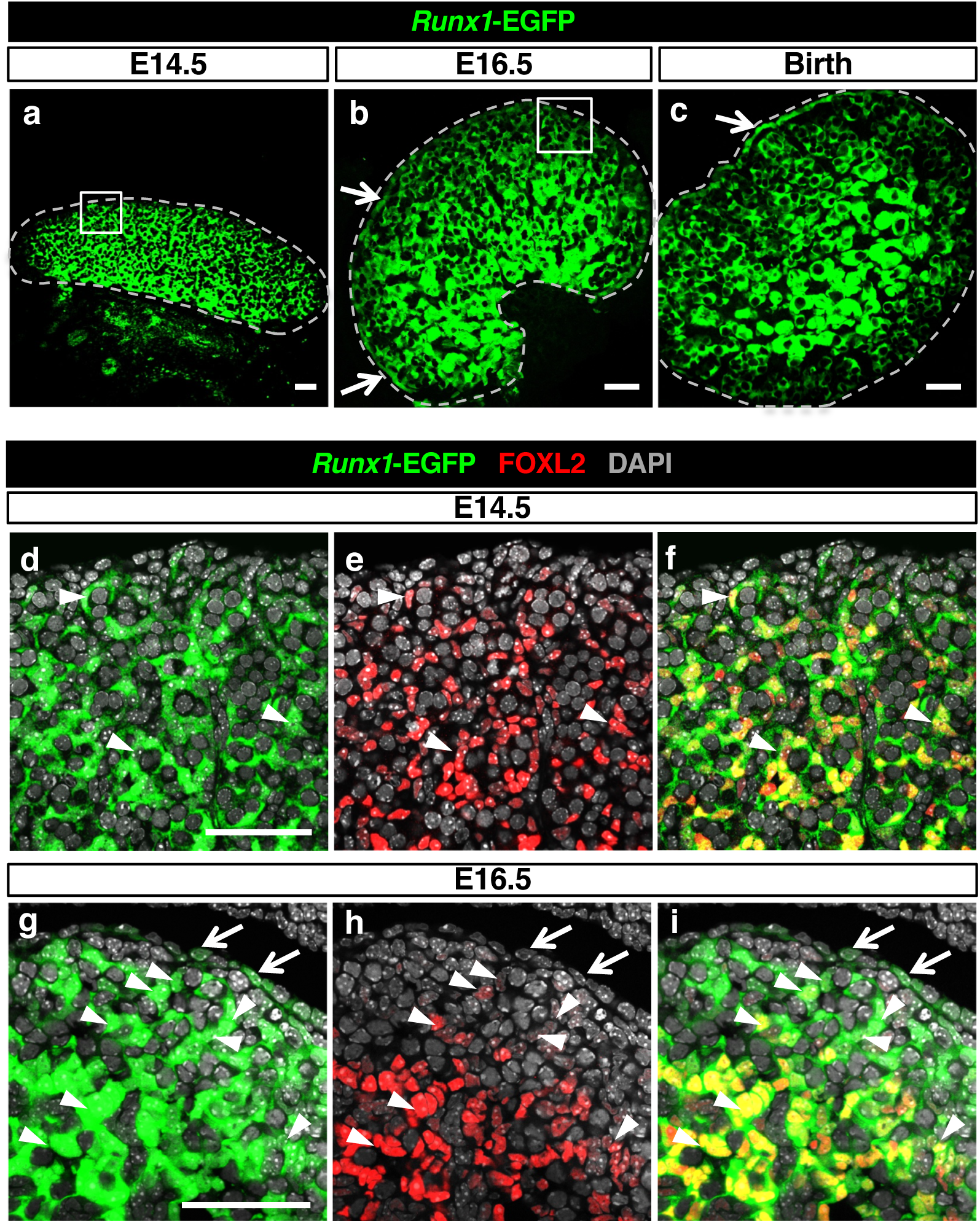
*Runx1* expression is maintained in granulosa cells throughout fetal ovarian development. **(a-c)** Immunofluorescence for EGFP on Tg(*Runx1*-EGFP) ovary sections at E14.5, E16.5 and birth. Scale bars: 50 μm. **(d-i)** Immunofluorescence for EGFP and the granulosa cell marker FOXL2 at E14.5 (d-f) and E16.5 (g-i), corresponding to the white square outlined areas in (a) and (b) respectively. *Runx1* is expressed in granulosa cells throughout ovarian development and the surface epithelium after E14.5. Dotted lines outline the gonads. Arrows: EGFP+ ovarian surface epithelium. Arrowheads: EGFP/FOXL2 double positive cells. Scale bars: 50 μm.

### Loss of *Runx1* leads to ovarian transcriptomic changes resembling those of ***Foxl2* knockout ovary**

The pre-granulosa cell-specific pattern suggests that RUNX1, a factor involved in cell lineage determination^25^, could contribute to granulosa cell differentiation and ovarian development. To investigate its specific role in gonadal somatic cells and avoid early embryonic lethality as a result of global deletion of *Runx1*^31, 32^, we generated a conditional knockout mouse model, in which *Runx1* was ablated in the SF1+ gonadal somatic cells^33^ (Fig. 4). While *Runx1* expression was ablated successfully in the fetal ovary (Fig. 4a), ovarian morphogenesis appeared normal at birth (Fig. 4b). The knockout (KO) ovary maintained its typical shape, with FOXL2+ pre-granulosa cells scattered throughout the ovary and TRA98+ germ cells located mostly in the cortex (Fig. 4b). Despite its normal morphology, *Runx1* KO newborn ovaries exhibited aberrant transcriptomic profile: expression of 317 genes was altered significantly compared to the control (Fig. 4c; Dataset S1). The transcriptomic changes of *Runx1* KO ovary were reminiscent of the ovary lacking *Foxl2*, a conserved gene involved in ovarian differentiation/maintenance in vertebrates. In the mouse, loss of *Foxl2* results in normal ovarian morphogenesis at birth despite aberrant ovarian transcriptome^5^. When comparing the genes changed significantly in the *Runx1* KO (317 genes) with those affected by the loss of *Foxl2* (749 genes) in newborn ovary, we found that 41% of the genes differentially expressed in *Runx1* KO (129/317) were also misregulated in the absence of *Foxl2* (Fig. 4c). The large majority of these 129 genes (93%; 120 genes) were similarly changed by the loss of either *Runx1* or *Foxl2*: 69% were downregulated in both KOs (89 genes) and 24% were upregulated in both KOs (31 genes; Dataset S1). One possible explanation for these common transcriptomic changes in *Runx1* and *Foxl2* KOs is that *Runx1* could be part of the same signaling cascade as *Foxl2*. However, analysis of *Runx1* expression in *Foxl2* KO newborn ovaries did not detect any changes in *Runx1* expression in the absence of *Foxl2* (Fig. 4d). Conversely, *Foxl2* expression was not changed in the absence of *Runx1*, indicating that *Runx1* and *Foxl2* are regulated independently of each other in the fetal ovary.

**Figure 4:**
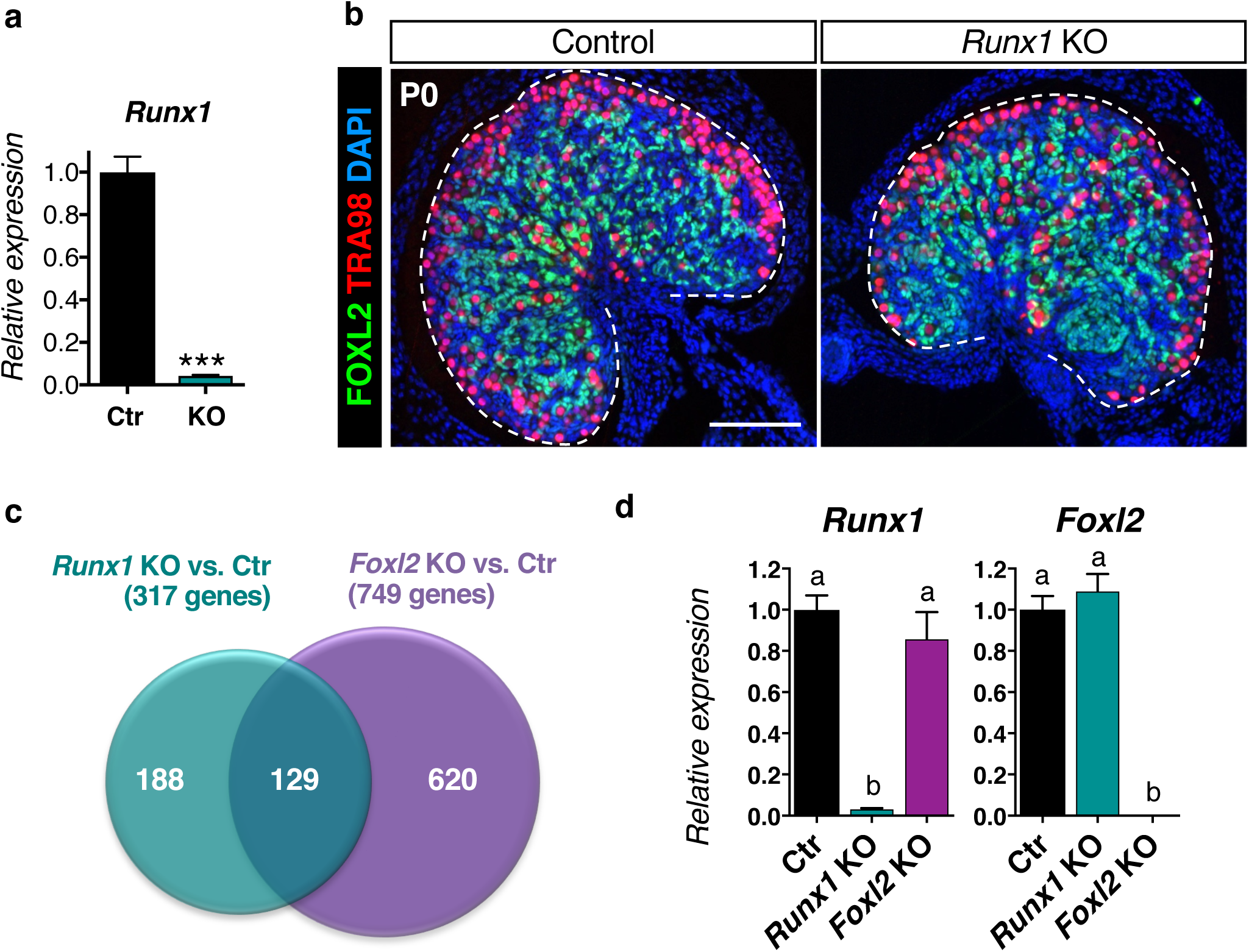
*Runx1* KO newborn ovaries present normal morphogenesis but share common transcriptomic changes with *Foxl2* KO ovaries. **(a)** Validation of *Runx1* knockout in E14.5 fetal ovaries by quantitative PCR (n=5); unpaired Student’s t-test \*\*\**P*<0.001. Values are presented as mean ± s.e.m. **(b)** Immunofluorescence for granulosa cell marker FOXL2, germ cell marker TRA98 and nuclear counterstain DAPI (blue) in control and *Runx1* KO ovaries at birth (P0). Scale bar: 100 μm. **(c)** Venn diagram comparing the 317 genes differentially expressed in newborn *Runx1* KO vs. Control ovaries (green circle) with the 749 genes differentially expressed in newborn *Foxl2* KO vs. Control ovaries (purple circle). Genes differentially expressed were identified by microarray (n=4/genotype; fold change >1.5, p<0.05). **(d)** Quantitative PCR analysis of *Runx1* and *Foxl2* mRNA expression in control, *Runx1* KO, and *Foxl2* KO newborn ovaries (n=5/genotype). Values are presented as mean ± s.e.m. One-way ANOVA, *P* < 0.05. Bars with different letters (a, b) are significantly different.

### Inactivation of both *Runx1* and *Foxl2* results in masculinization of the fetal ovaries

The common transcriptomic changes identified in *Runx1* and *Foxl2* KO ovaries raised the question whether RUNX1 and FOXL2 could play redundant or complementary roles in supporting cell differentiation. We therefore generated *Runx1/Foxl2* double KO mice (hereafter referred as DKO) and compared ovarian differentiation in the absence of *Runx1, Foxl2*, or both (Fig. 5 and S2-3). At E15.5, differentiation of supporting cells into Sertoli cells in the testis or pre-granulosa cells in the ovary has already been established. For instance, the transcription factor DMRT1, which is involved in the maintenance of Sertoli cell identity^18^, is expressed in Sertoli cells but not pre-granulosa cells (Fig. 5a & e). At this stage, DMRT1 is also present in a few germ cells in both testis and ovary^34^. Similar to the control ovaries, ovaries lacking either *Runx1* or *Foxl2* had no DMRT1 proteins in the supporting cells (Fig. 5a-c). However, the combined loss of *Runx1* and *Foxl2* resulted in aberrant expression of DMRT1 in the supporting cells of the fetal ovary (Fig. 5d). At birth, *Runx1/Foxl2* DKO ovary formed structures similar to fetal testis cords in the center, with DMRT1+ cells surrounding clusters of germ cells (Fig. 5i). Such structure was not observed in *Runx1* or *Foxl2* single KO ovaries with the exception that DMRT1 protein started to appear in a few supporting cells in the newborn *Foxl2* KO ovaries, in what appears to be one of the first signs of postnatal masculinization of *Foxl2* KO ovaries at the protein level (Fig. 5h). Contrary to DMRT1, SOX9 protein, a key driver of Sertoli cell differentiation^35, 36^, was not detected in the *Runx1/Foxl2* DKO newborn ovaries (Fig. 6). Our results demonstrate that a combined loss of *Runx1* and *Foxl2* induces partial masculinization of the supporting cells during fetal development of the ovary.

**Figure 5:**
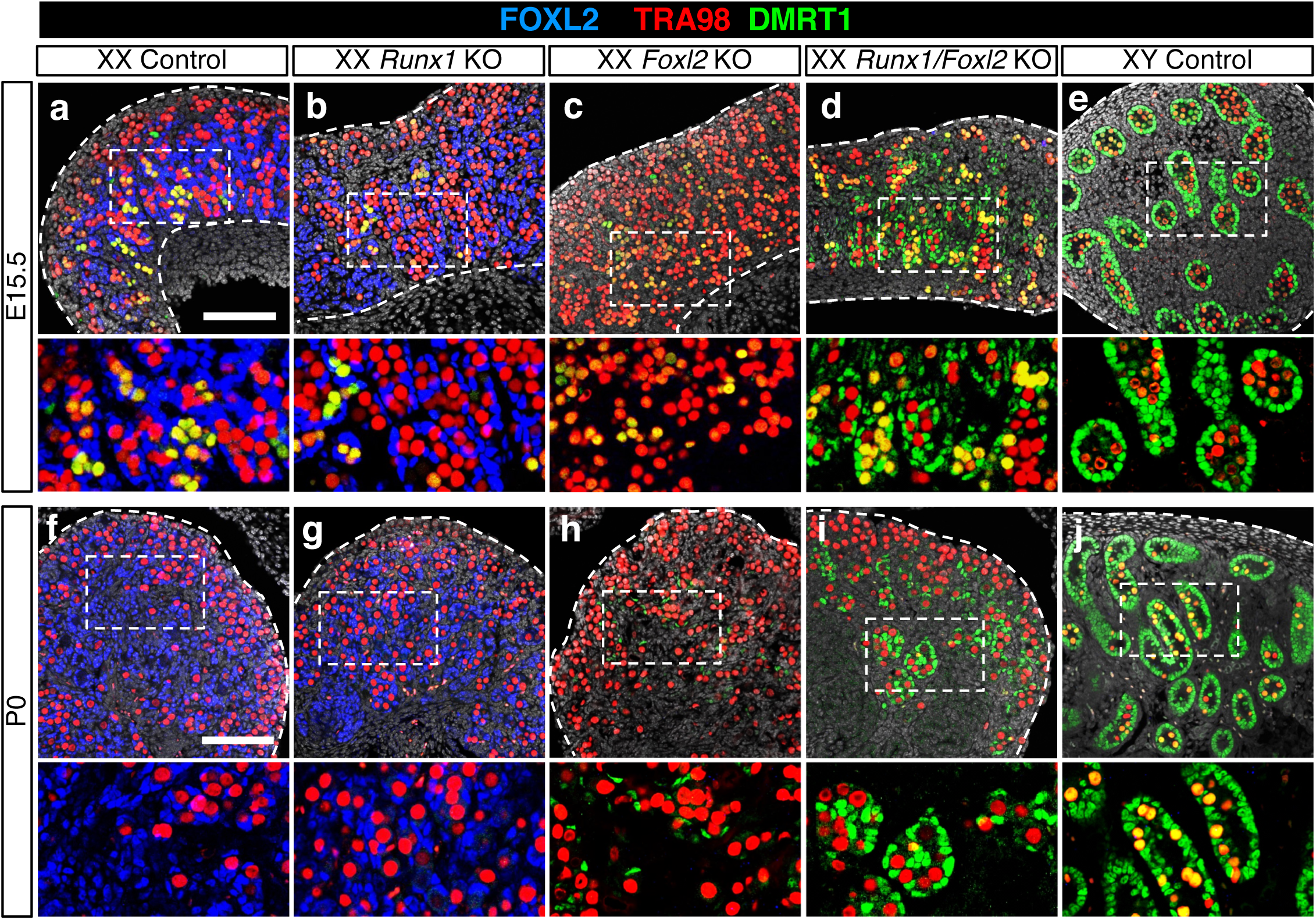
Combined loss of *Runx1* and *Foxl2* results in masculinization of the fetal ovaries. **(a-j)** Immunofluorescence for the Sertoli cell and germ cell marker DMRT1, the germ cell marker TRA98 and the granulosa cell marker FOXL2 in control, *Runx1* KO, *Foxl2* KO, *Runx1/Foxl2* double knockout ovaries and control testes at E15.5 **(a-e)** and birth **(f-j)**. The grey represents DAPI nuclear staining. Dotted lines outline the gonads. Higher magnifications are shown for the outlined boxes in **a-e** and **f-j** respectively. Scale bars: 100 μm.

**Figure 6:**
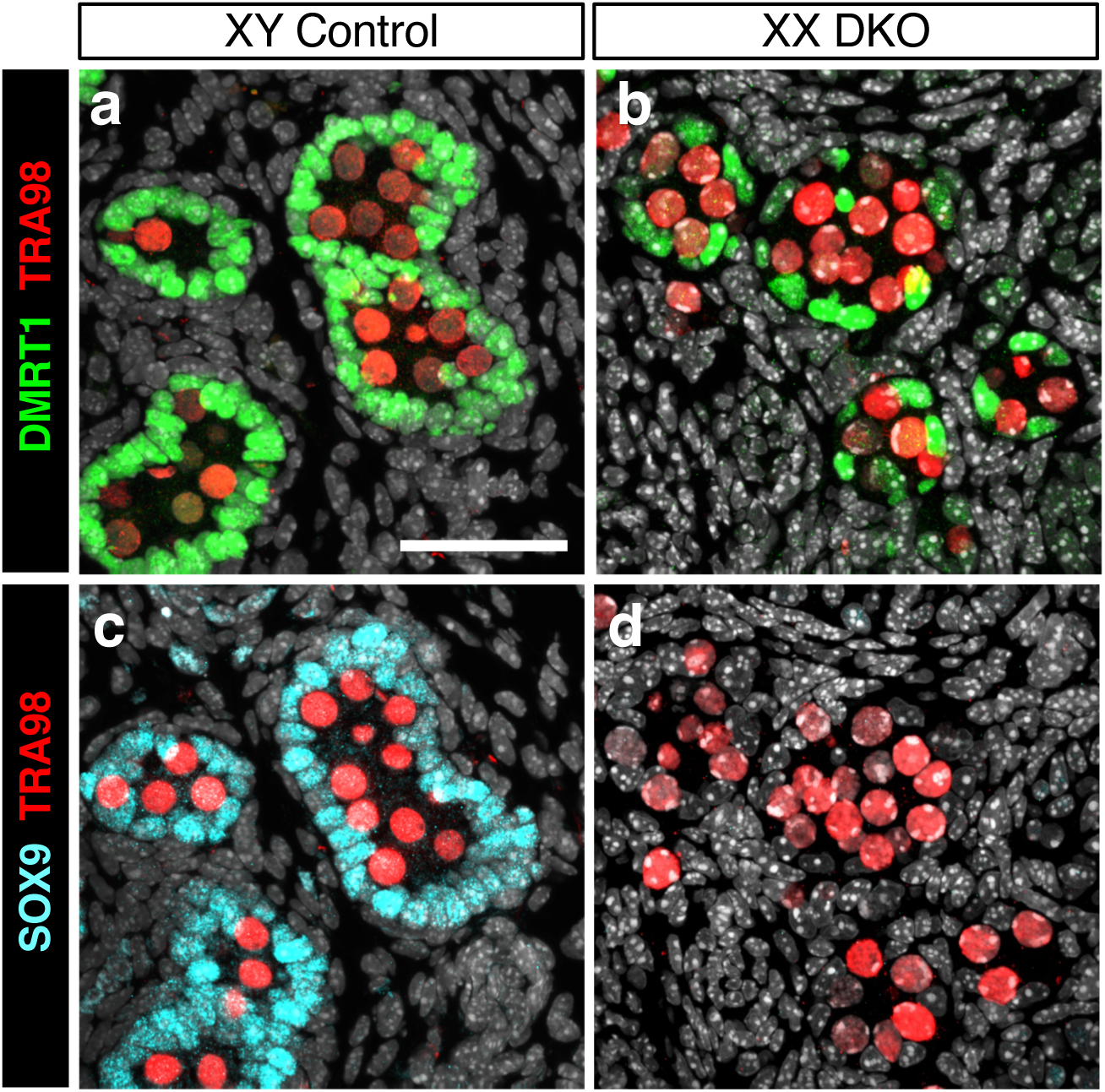
SOX9 protein is not detected in *Runx1*/*Foxl2* DKO newborn ovaries. Immunofluorescence for Sertoli cell markers DMRT1 and SOX9 and germ cell marker TRA98 on consecutive sections in control testis (a & c) and *Runx1/Foxl2* DKO ovary (b & d) at birth. Scale bar: 50 μm.

To further characterize the impacts of the combined loss of *Runx1* and *Foxl2* on ovarian differentiation, we compared the transcriptome of *Runx1/Foxl2* DKO newborn ovaries with the transcriptomes of control, *Runx1*, or *Foxl2* single KO ovaries (Fig. 7 and Dataset S2). Between *Runx1/Foxl2* DKO and control ovaries, 918 genes were differentially expressed, with 499 genes downregulated and 419 genes upregulated in the DKO ovary (fold-change >1.5; p<0.05; Dataset S3). The heat-map for the 918 differentially expressed genes in *Runx1/Foxl2* DKO ovaries demonstrated allele-specific impacts with a mild and often non-significant effect in the absence of *Runx1*, an intermediate/strong effect in the absence of *Foxl2*, and a strongest effect in the absence of both *Runx1* and *Foxl2* (Fig. 7a). Gene ontology (GO) analysis revealed that the downregulated genes were associated with “ovarian follicle development” and “female gonad development” whereas “male sex determination” was the most significantly enriched process for the upregulated genes (Fig. S4). To determine the contribution of *Runx1* and *Foxl2* to the transcriptomic changes, the genes significantly downregulated (Fig. 7b, d & e; Dataset S4) or upregulated (Fig. 7c, f, & g; Dataset S4) were compared among *Runx1/Foxl2* single and double knockouts. Conforming to the hierarchical clustering (Fig. 7a), *Foxl2* was the main contributor to the transcriptional changes observed in *Runx1/Foxl2* DKO: 61% of the genes downregulated in DKO were also downregulated in *Foxl2* KO (304/499 genes in the overlapping region between purple and red circle in Fig. 7b) and 43% of the genes upregulated in DKO were also upregulated in *Foxl2* KO (182/419 genes in the overlapping region between purple and red circle in Fig. 7c). In addition, some genes appeared to be controlled by both *Foxl2* and *Runx1*, and were significantly downregulated (66 genes in the overlapping region between the three circles in Fig. 7b) or upregulated (29 genes in the overlapping region between the three circles in Fig. 7c) in all three knockouts. For instance, the genes *Fst* and *Cyp19a*, both involved in granulosa cell differentiation/function ^37, 38^, were downregulated in *Runx1* KO, *Foxl2* KO, and more repressed in *Runx1/Foxl2* DKO (Fig. 7d). On the other hand, desert hedgehog (*Dhh*) was upregulated in all 3 knockouts, with highest expression in the DKO (Fig. 7f). Notably, 30% of the genes downregulated (151/499 genes; Fig. 7b) and 52% of the genes upregulated (219/419 genes; Fig. 7c) in *Runx1/Foxl2* DKO were not significantly changed in *Runx1* or *Foxl2* single KO. Most of the genes in this category were mildly changed in the single knockouts without reaching the cut-off of the microarray analysis. For instance, *Foxp1*, a gene whose expression is enriched in pre-granulosa cells^39^, was significantly downregulated only in *Runx1/Foxl2* DKO (Fig. 7e) whereas *Fgf9*, a Sertoli gene contributing to testis differentiation^40^ and *Pdgfc* were significantly upregulated only in *Runx1/Foxl2* DKO (Fig. 7g).

**Figure 7:**
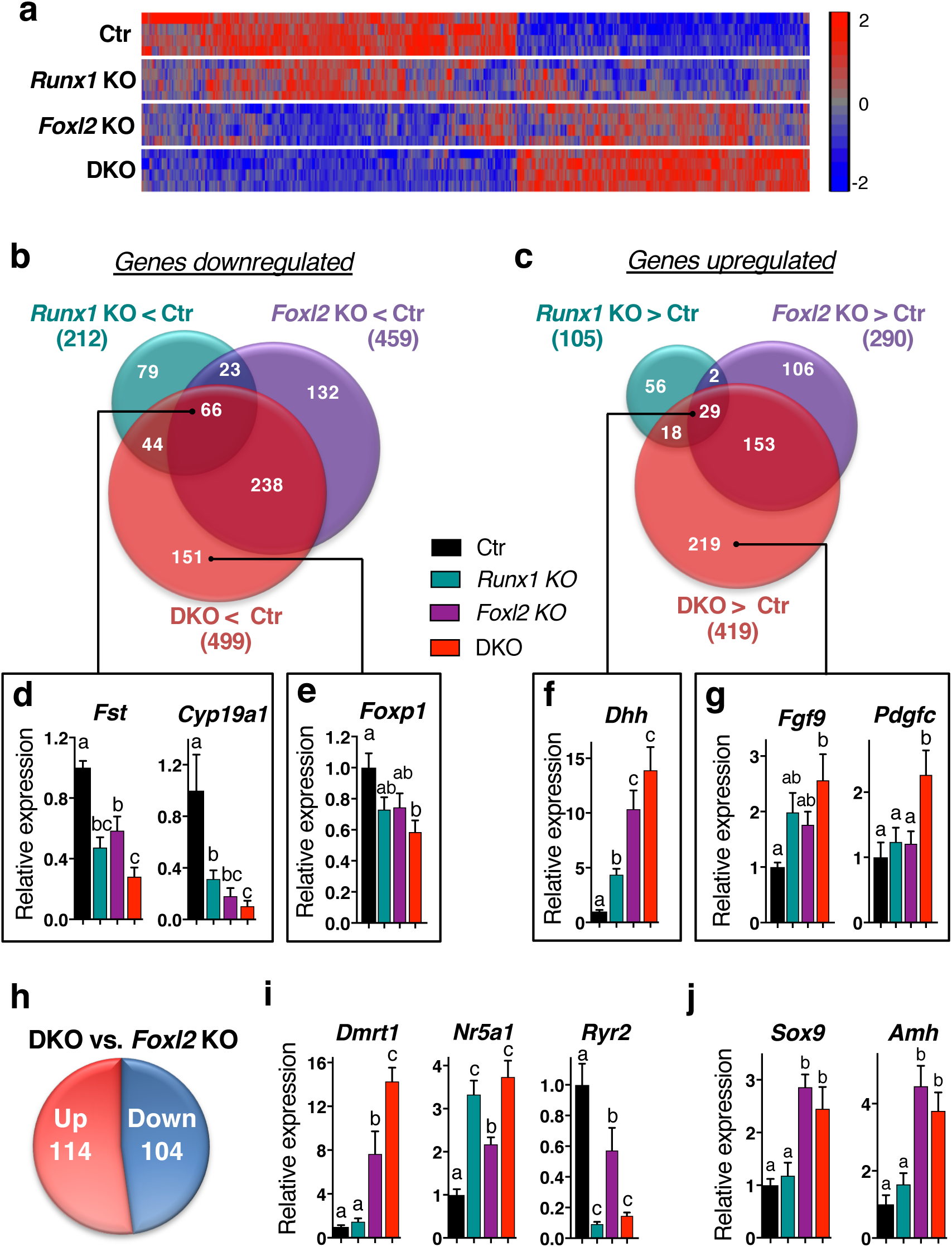
Comparison of *Runx1* KO, *Foxl2* KO and *Runx1/Foxl2* double KO transcriptomes. **(a)** Heat-map for the 918 genes differentially expressed in *Runx1/Foxl2* double knockout (DKO) vs. control (Ctr) newborn ovaries. The heat map shows the expression of these 918 genes in control, *Runx1* KO, *Foxl2* KO and *Runx1/Foxl2* DKO ovaries (microarray; n=4/genotype; One-way ANOVA; fold change >1.5, *P*<0.05). **(b-c)** Venn diagram comparing the genes downregulated (**b**) or upregulated (**c**) in *Runx1* KO (green circle), *Foxl2* KO (purple circle) and *Runx1/Foxl2* DKO (purple circle) newborn ovaries. **(d-g)** Validation by quantitative PCR of genes identified in the Venn diagrams as significantly downregulated in all three KO **(d),** or only in *Runx1/Foxl2* DKO **(e)**, or significantly upregulated in all in all three KO **(f)** or only in *Runx1/Foxl2* DKO **(g)**. **(h-i)** Identification of the genes differentially expressed in *Runx1/Foxl2* DKO vs. *Foxl2* KO ovaries and validation of candidate genes by quantitative PCR. **(j)** Expression of *Sox9* and *Amh* is not changed in *Runx1/Foxl2* DKO compared to *Foxl2* KO ovaries. For all the qPCR data, values are presented as mean ± s.e.m (n=5/genotype). One-way ANOVA, *P* < 0.05. Bars with different letters (a, b, c) are significantly different.

Comparing to *Foxl2* single KO female (Fig. 5), in which sex reversal only became apparent postnatally^5^, *Runx1/Foxl2* DKO female exhibited masculinization of the ovaries with visible morphological changes before birth. To determine how the loss of *Runx1* contributed to the early masculinization of *Runx1/Foxl2* DKO ovaries, we identified the potential RUNX1-regulated genes by comparing the transcriptomes of *Runx1/Foxl2* DKO and *Foxl2* single KO ovaries. A total of 218 genes were differentially expressed between *Runx1/Foxl2* DKO and *Foxl2* KO ovaries, with 114 genes significantly upregulated, and 104 genes significantly downregulated in the DKO (fold-change >1.5; p<0.05; Fig. 7h and Dataset S5). Expression of most of these genes was already altered in *Foxl2* single KO ovaries; however, the additional loss of *Runx1* exacerbated their mis-expression. For instance, the pro-testis gene *Dmrt1* and *Nr5a1* were significantly upregulated whereas the granulosa-cell enriched transcripts *Fst* and *Ryr2* were further downregulated at birth (Fig. 7d & i). On the other hand, the additional loss of *Runx1* did not cause further upregulation of the Sertoli genes *Sox9* and *Amh* at birth, suggesting that *Runx1* does not contribute to their repression in the ovary (Fig. 7j). Overall, the transcriptomic analyses of *Runx1/Foxl2* single and double knockouts reveal that the additional loss of *Runx1* amplifies the mis-expression of genes already altered by the sole loss of *Foxl2*.

### RUNX1 shares genome-wide chromatin occupancy with FOXL2 in the fetal ovary

The masculinization of *Runx1/Foxl2* DKO fetal ovaries and the transcriptomic comparisons of *Runx1/Foxl2* single and double KO ovaries suggest some interplay between RUNX1 and FOXL2 to control granulosa cell identity. The fact that RUNX1 and FOXL2 are both transcription factors raised the question whether this interplay could occur at the chromatin level. We have previously identified the chromatin occupancy of FOXL2 during ovarian differentiation by chromatin immunoprecipitation followed by whole genome sequencing or ChIP-seq^41^. We performed additional *de novo* motif analyses on the genomic regions bound by FOXL2 in the fetal ovary, and discovered that several other DNA motifs were co-enriched with the FOXL2 DNA motif (Fig. 8a). Among them, RUNX DNA motif was the second most significantly co-enriched motif. The other motifs were for CTCF, a factor involved in transcriptional regulation, enhancer insulation and chromatin architecture^42^, and for the DNA motif recognized by multiple members of the nuclear receptor family including liver receptor homolog-1 (LRH1 encoded by *Nr5a2*) and SF1 (encoded by *Nr5a1*), a known co-factor of FOXL2^43, 44^. DNA motifs for TEAD transcription factors of the Hippo pathway, ETS, NFYA and GATA4 were also significantly enriched. The enrichment of RUNX motif with FOXL2 binding motif suggests that RUNX1, the only RUNX enriched in pre-granulosa cells, could bind similar genomic regions to FOXL2 in the fetal ovary. To confirm this hypothesis, we performed ChIP-seq for RUNX1 in E14.5 ovaries (Dataset S6), the same stage the FOXL2 ChIP-seq data were obtained^41^. The top *de novo* motif identified in RUNX1 ChIP-seq (p<1e-559) matched the RUNX motif^45^ (Fig. 8b), and corresponded to the motif that was co-enriched with FOXL2 in FOXL2 ChIP-seq (Fig. 8a). A total of 10,494 RUNX1 binding peaks were identified in the fetal ovary, with the majority of the peaks located either in the gene body (Fig. 8c; 25% exon and 22% intron), or close upstream of the transcription start site TSS (30% Promoter: <1kb of TSS; 12% Upstream: −10 to −1kb of TSS). Comparison of genome-wide chromatin binding of RUNX1 and FOXL2 in the fetal ovary revealed significant overlap: 54% (5,619/10,494) of RUNX1 peaks overlapped with FOXL2 peaks (Fig. 8d).

**Figure 8:**
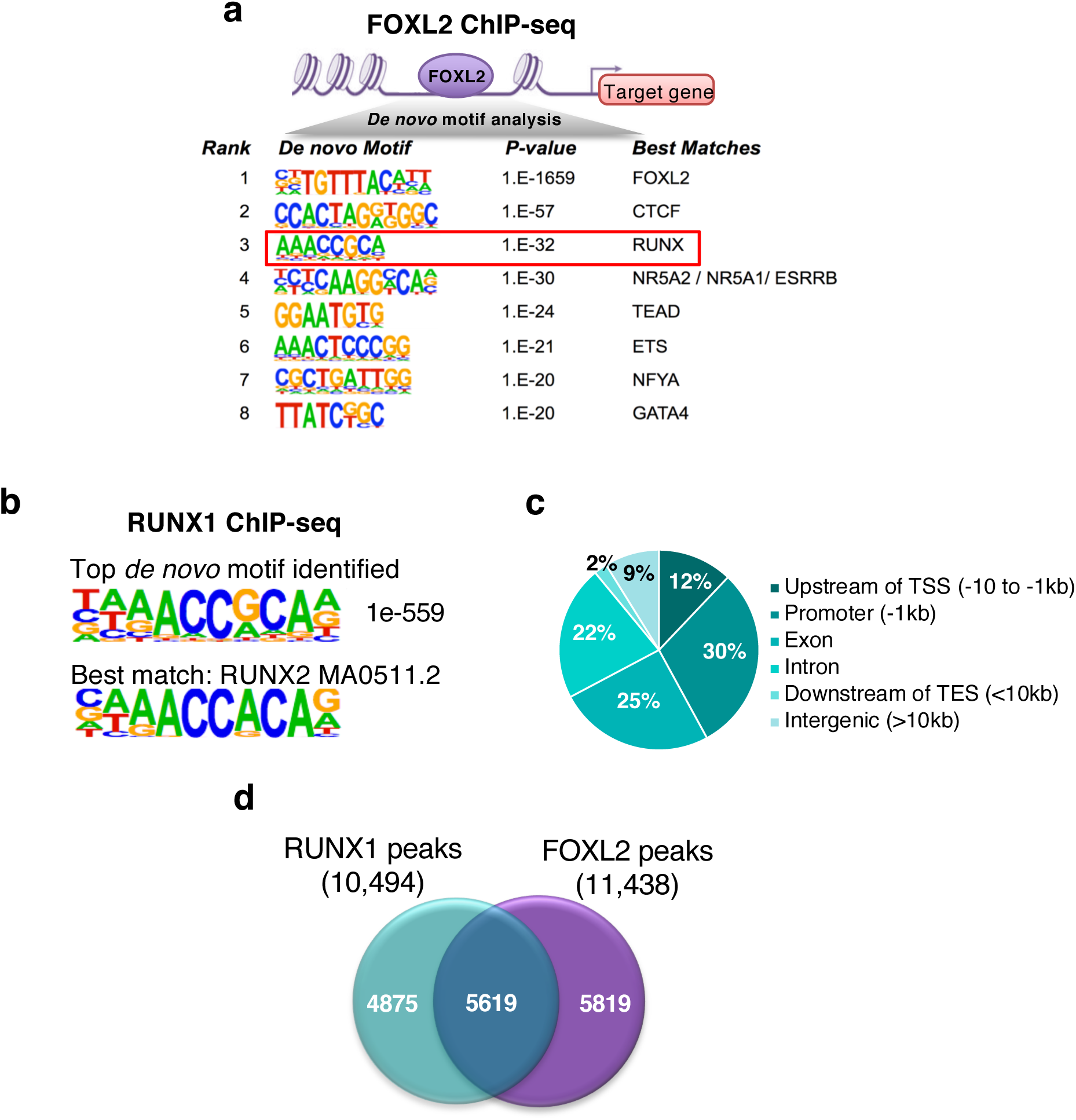
RUNX1 and FOXL2 exhibit extensive overlaps in chromatin binding in fetal ovaries. **(a)** *de novo* motif analysis of FOXL2 peaks identifies enrichment of RUNX motif along with FOXL2 motif in E14.5 ovaries. **(b)** The top *de novo* motif for RUNX1 ChIP-seq in E14.5 ovaries corresponds to a RUNX motif. **(c)** Distribution of genomic location of the 10,494 RUNX1 binding peaks. TSS: Transcription Start Site; TES: Transcription End Site. **(d)** Comparison of RUNX1 (10,494 peaks) and FOXL2 (11,438 peaks) chromatin occupancy in E14.5 ovaries.

The transcriptomic data from *Runx1/Foxl2* DKO ovaries provided us a list of genes significantly changed as a result of the absence of *Runx1*, *Foxl2*, or both (Fig. 7). To identify potential direct target genes of RUNX1 or/and FOXL2, we focused on the 918 genes differentially expressed in *Runx1/Foxl2* DKO ovaries, and determined which genes were nearest to RUNX1 or/and FOXL2 binding peaks (Fig. 9a and Dataset S7). More than 50% of these genes (492/918; Fig. 9a) were the closest gene to RUNX1 or/and FOXL2 peak. Some of these genes were nearest to only FOXL2 peaks (116 genes in Fig. 9a). For example, *Pla2r1*, a transcript enriched in pre-granulosa cells^39^ and similarly downregulated in both *Foxl2* KO and *Runx1/Foxl2* DKO fetal ovaries (Fig. 9b), contained two FOXL2-specific peaks, one in the promoter and one in the first intron. On the other hand, 102 genes (Fig. 9a) had RUNX1 specific peaks near their genomic locations. For instance, *Ryr2*, another transcript enriched in pre-granulosa cells^39^, was strongly downregulated in both *Runx1* KO and *Runx1/Foxl2* DKO fetal and newborn ovaries (Fig. 7i and Fig. 9c), and contained one RUNX1 specific peak in its intronic region. Finally, 274 genes were the closest genes to peaks for both RUNX1 and FOXL2, with the majority of them (197 genes) nearest to overlapping peaks for RUNX1 and FOXL2 (Fig. 9a). Most of these genes (74%; 146/197 genes) were downregulated in *Runx1/Foxl2* DKO ovaries (Dataset S7). For instance, the granulosa cell enriched genes *Fst* and *Itpr2*, both downregulated in *Runx1/Foxl2* single and double KO ovaries (Fig. 7d and 9d), contained common binding peaks for FOXL2 and RUNX1 (Fig. 9a and 9d). For *Fst*, this binding of RUNX1 and FOXL2 was located in its first intron, in the previously identified regulatory region that contributes to its expression^41, 46^. On the other hand, the Sertoli cell gene *Dmrt1*, strongly upregulated in *Runx1/Foxl2* DKO (Fig. 5 and 7i), contained a common binding site for FOXL2 and RUNX1 near its promoter (Fig. 9a). Taken together, our results reveal that RUNX1, a transcription factor expressed in the fetal ovary of various vertebrate species, contributes to ovarian differentiation and maintenance of pre-granulosa cell identity through an interplay with FOXL2 that occurs at the chromatin level.

**Figure 9:**
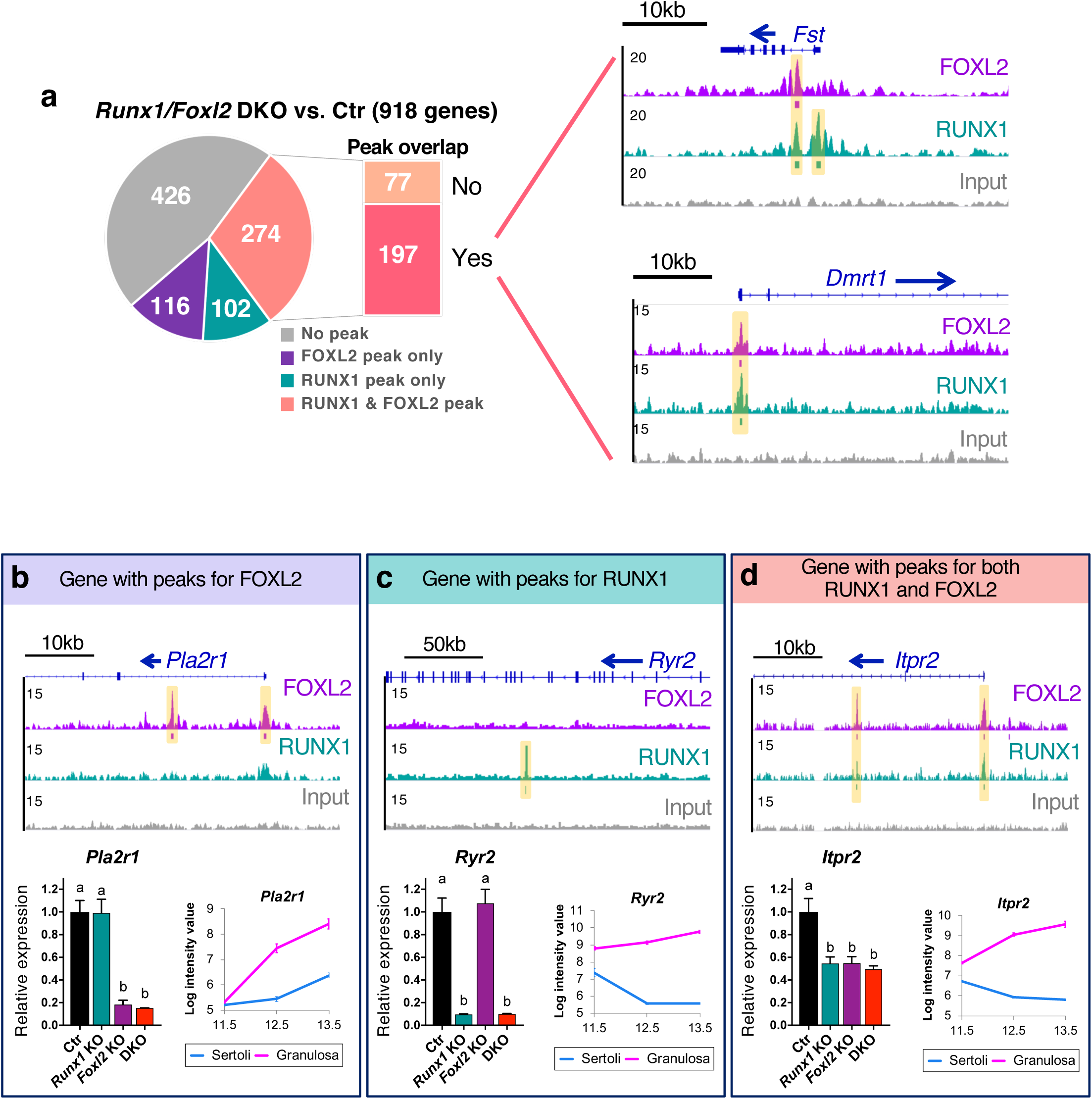
Identification of the genes significantly changed in Runx1/Foxl2 DKO that are nearest to RUNX1 and/or FOXL2 genomic binding peaks. **(a)** Pie-chart of the genes significantly changed in *Runx1/Foxl2* DKO based on the presence of peak for FOXL2 and/or RUNX1. Genome browser view of 2 key genes significantly changed in *Runx1/Foxl2* DKO ovaries and bound by RUNX1 and FOXL2 in E14.5 ovaries. Blue arrows: gene orientation; Orange highlighted area: significant binding peaks identified by HOMER. **(b-d)** Examples of genes affected in *Runx1/Foxl2* DKO ovary and bound by FOXL2 or/and RUNX1. For each gene, we show the genome browser view of RUNX1 and/or FOXL2 binding in E14.5 ovaries, the gene expression by quantitative PCR in *Runx1/Foxl2* single and double knockouts at E15.5 (n=4/genotype; mean ± s.e.m.; One-way ANOVA, P < 0.05.), and the gene expression in fetal Sertoli and granulosa cells from E11.5 to E13.5^39^. Bars with different letters (a, b) are significantly different.

## Discussion

### RUNX1 is a part of the multi-component network that controls pre-granulosa cell identity

The molecular events that specify granulosa cell fate are complex and non-linear, involving several signaling pathways that have redundant and complementary functions. This is in contrast to the fetal testis, where the molecular pathway driving its differentiation appears linear and sequential. Removal of one of the top pieces in the testis differentiation pathway has a domino effect that prevents induction of downstream events. This is exemplified by the complete gonadal sex-reversal in gain- or loss-of-function mouse models for SRY or SOX9, the two transcription factors responsible for the initiation of the testis morphogenesis^28, 35, 47^. This is not the case in the mouse ovary, where no single-gene loss/mutation results in a complete ovary-to-testis sex-reversal. For instance, defects in the WNT4/RSPO1/beta-catenin or loss of *Foxl2* causes a late or postnatal partial ovary-to-testis sex-reversal, while the combined loss of *Foxl2* and elements of the WNT4/RSPO1/beta-catenin pathway (*Wnt4* or *Rspo1*) leads to ovary-to-testis sex-reversal more pronounced than each single knockout model in the mouse^8, 9^. In this study, we demonstrated that *Runx1* contributes to the molecular network controlling pre-granulosa cell differentiation. Loss of *Runx1* in somatic cells of the ovaries altered ovarian transcriptome but did not affect ovarian morphogenesis at birth. In contrast, the combined loss of *Runx1* and *Foxl2* compromised pre-granulosa cells identity. Loss of *Runx1* or *Foxl2* affected a common set of genes, and these transcriptomic changes were enhanced in the absence of both genes, reaching a threshold that masculinized the fetal ovary. One of the most striking changes was the expression of DMRT1 in the fetal supporting cells. DMRT1 is a key driver of Sertoli cell differentiation and testis development in various species^15, 20^. In the fly, *doublesex* (*dsx*), an ortholog of mammalian *DMRT1*, controls testis differentiation^15^. Intriguingly, *runt*, the fly ortholog of *RUNX1*, drives ovarian differentiation by antagonizing the testis-specific transcriptional regulation of *dsx*^21^. In the mouse, testis differentiation is not controlled by DMRT1 but by SOX transcriptions factors SRY and its direct target SOX9. However, RUNX1 does not appear to contribute to the repression of the key pro-testis gene *Sox9* in the fetal ovary and SOX9 protein was not detected in *Runx1/Foxl2* DKO ovary at birth. This is in contrast with the phenotype of *Wnt4/Foxl2* DKO newborn ovaries where SOX9 was strongly upregulated, and as a consequence the ovaries were more masculinized^9^. Overall, our findings suggest that slightly different pro-ovarian networks control the repression of the evolutionary conserved pro-testis genes *Sox9* and *Dmrt1*: *Sox9*, which plays a primary role in Sertoli cell differentiation in the mouse, is repressed by an interplay between the WNT4/RSPO1/beta-catenin and FOXL2^8, 9^. On the other hand, *Dmrt1*, which has taken a secondary role in Sertoli cell differentiation in the mouse, is repressed by an interplay between RUNX1 and FOXL2.

Seeking the mechanisms underlying the interplay between RUNX1 and FOXL2, we identified that RUNX DNA binding motif is significantly co-enriched with FOXL2 motif in genomic regions bound by FOXL2 in the fetal ovary. Moreover, RUNX1 genome-wide occupancy partially overlaps with FOXL2 in the fetal ovary, suggesting that they bind common regulatory regions to control granulosa-cell identity and ovarian development. By themselves, RUNX proteins are weak transcription factors, and they require other transcriptional regulators to function as either repressors or activators of transcription^26^. Interplay between RUNX1 and several members of the forkhead transcription factor family has been documented in different tissues. For instance, RUNX1 is a co-activator of FOXO3 in hepatic cells^48^. Similarly, an interplay between RUNX1 and FOXO1/FOXO3 was demonstrated in breast epithelial cells where a subset of FOXO target genes were jointly regulated with RUNX1^49^. Another forkhead protein, FOXP3, acts with RUNX1 to control gene expression in T-cells^50^ and breast epithelial cells^51^. Such cooperation in various tissues suggest that the interplay between RUNXs and forkhead transcription factors maybe an evolutionary conserved phenomenon. In addition to the genes co-regulated with FOXL2, we identified genes that were specifically mis-expressed in the absence of *Runx1* but not *Foxl2*. Genome-wide analyses of RUNX1 binding in the fetal ovary also identified genomic regions bound by RUNX1 but not FOXL2. These results suggest that RUNX1 could also contribute to ovarian development or function independently of FOXL2.

### RUNX1, a marker of gonadal supporting cell identity

RUNX1 contributes to cell fate determination in various developmental processes such as hematopoiesis and hair follicle development. Depending on its interplay with other signal transduction pathways or co-factors, RUNX1 controls which path the precursor cells take when they are at the crossroad between cell proliferation/renewal and lineage-specific commitment^25^. We discovered that *Runx1* has an ovary-biased expression during gonad differentiation in various vertebrate species, including turtle, rainbow trout, goat, mouse, and human. In the mouse embryonic gonads, *Runx1* is first detected in the supporting cells in a non-sexually dimorphic way at the onset of sex determination. While its expression is maintained in the ovary, *Runx1* appears to be actively repressed in the testis between E11.5 and E12.5 as the supporting cells commit to Sertoli cell fate. The suppression of *Runx1* in the fetal testis is corroborated by previously published data from a time-course transcriptomic analysis during early gonad development^52^ and single-cell sequencing analysis of SF1+ progenitor cells^53^. The time-course of Sertoli cell differentiation at the single-cell level revealed that *Runx1* follows an identical spatiotemporal pattern of expression with *Sry^53^*. In the mouse, *Sry* expression in Sertoli cells is quickly turned off after the initiation of testis differentiation, and it is suspected that the repression of *Sry* is due to a negative feedback loop by downstream pro-testis genes. The similar pattern of downregulation of *Runx1* in the testis after E11.5 raises the possibility that *Runx1* is downregulated by a similar signaling pathway. Regulation of *Runx1* gene expression is complex, and several enhancers that confer tissue-specific expression have been identified^54^. It remains to be determined how *Runx1* expression is controlled in the gonads, and how it is actively repressed in the fetal testis.

In contrary to the testis, the fetal ovary maintains expression of *Runx1* in the supporting cells as they differentiate into granulosa cells. During ovarian differentiation, granulosa cells arise from two different waves at different stages of development: the first cohort of granulosa cells arises from the bipotential supporting cell precursors that are able to differentiate into either Sertoli cells or pre-granulosa cells during sex determination^55^. The second wave of granulosa cells that eventually populate the cortical region of the ovary appears later in gestation. These cells of the second wave arise from LGR5+ cells of the ovarian surface epithelium that ingress into the ovary from E15.5 to postnatal day 4 and eventually become LGR5-/FOXL2+ granulosa cells ^29, 30, 56^. This timing of the establishment of the second cohort of granulosa cells correlates with the expression of *Runx1*-EGFP in a subset of cells in the surface epithelium and in granulosa cells of the ovarian cortex at E16.5 and birth. These results suggest that *Runx1* marks granulosa cell precursors that will give rise to the second wave of FOXL2+ granulosa cells in the cortex. Therefore, both expression at onset of sex determination and at the surface epithelium/cortex at the time of the second wave of granulosa cells recruitment suggest that *Runx1* is activated in cells that are primed to become supporting/granulosa cells.

### Towards the identification of granulosa cell genomic signatures

Multiple transcription factors, rather than a single one, often form complex genetic regulatory networks that control cell fate determination. Genomic sequence motifs or cis-regulatory elements for Sertoli cells, the supporting cell lineage in the testis, were identified by combined analyses of SOX9 and DMRT1 ChIP-seq and by motif prediction^57^. These “Sertoli cell signatures” are composed of organized binding motifs for transcription factors critical for Sertoli cell differentiation, including SOX9, GATA4 and DMRT1. These Sertoli cell signatures, present in mammals and other vertebrates, could represent a conserved regulatory code that governs the cascade of Sertoli cell differentiation, regardless whether it primarily relies on SOX transcription factors like SRY in mammals, or on DMRT1 like in several vertebrate species. Similarly, one would expect the presence of conserved “granulosa cell signature” genomic regions that confers granulosa cell differentiation. Since FOXL2 is a highly conserved gene in granulosa cell differentiation in vertebrates, we used FOXL2 as an anchor factor to identify other factors that could take part in the regulatory network controlling granulosa cell differentiation/function. Unbiased analyses of the motifs co-enriched with FOXL2 motif in the fetal ovary identified the RUNX motif as one of the most co-enriched motifs. In addition to the RUNX motif, motifs for CTCF, nuclear receptors SF-1/LRH-1/ESRRB and transcription factors TEADs and GATAs were also significantly enriched with FOXL2 consensus motif in FOXL2-bound chromatin regions. For many of these transcription factors, their potential role in gonad differentiation is unknown or limited. For example, the transcription factors of the TEAD family belong to the Hippo pathway, which is involved in the regulation of Sertoli cell gene expression in the fetal gonads^58^. However, the potential involvement of the hippo pathway in granulosa cell differentiation has not been investigated.

In conclusion, we identified RUNX1 as a transcription factor involved in pre-granulosa cell differentiation. RUNX1 delineates the supporting cell lineage and contributes to granulosa cell differentiation and maintenance of their identity through an interplay with FOXL2. Our findings provide new insights into the genomic control of granulosa cell differentiation, and pave the way for the identification of novel transcription factors and cis-signatures contributing to the fate determination of granulosa cells and the consequent formation of a functional ovary.

## Methods

### Mouse models

Tg(*Runx1*-EGFP) reporter mouse was purchased from MMRRC (MMRRC_010771- UCD), and CD-1 mice were purchased from Charles River (stock number 022). *Runx1*^+/-^ (B6.129S-*Runx1^tm1Spe^*/J) and *Runx1^f^*^/f^ (B6.129P2-*Runx1^tm1Tani^*/J) mice were purchased from the Jackson Laboratory (stock numbers 005669 and 008772, respectively). *Sf1*- Cre^Tg/Tg^ mice^33^ were provided by late Dr. Keith Parker and *Foxl2*^+/-^ mice^59^ by Dr. David Schlessinger (National Institute on Aging), respectively. *Runx1* KO mice (*Sf1Cre^+/Tg^; Runx1^f/-^*) were generated by crossing *Runx1^f^*^/f^ females with *Sf1-Cre^+/Tg^; Runx1^+/-^* males. Controls were *Sf1-Cre^+/+^; Runx1^+/f^* littermates. *Runx1/Foxl2* double knockout mice (*Sf1Cre^+/Tg^; Runx1^f/-^; Foxl2^-/-^*) were generated by crossing *Runx1^f/f^; Foxl2^+/-^* females with *Sf1Cre^+/Tg^; Runx1^+/-^; Foxl2^+/-^* males. This cross also generated the single knockouts for *Runx1* (*Sf1Cre^+/Tg^; Runx1^f/-^; Foxl2^+/+^)* and *Foxl2* (*Sf1Cre^+/+^; Runx1^f/+^; Foxl2^-/-^*), and control littermates (*Sf1-Cre^+/+^; Runx1^+/f^; Foxl2^+/+^*). Time-mating was set up by housing female mice with male mice overnight and the females were checked for the presence of vaginal plug the next morning. The day when the vaginal plug was detected was considered embryonic day E0.5. All experiments were performed on at least four animals for each genotype. All animal procedures were approved by the National Institutes of Health Animals Care and Use Committee, and were performed in accordance with an approved National Institute of Environmental Health Sciences animal study proposal.

### Immunofluorescences

For the Tg(*Runx1*-EGFP) mice, gonads were collected and fixed in 4% paraformaldehyde for 1-2h at room temperature. Immunofluorescence experiments were performed on whole gonads at E11.5 and E12.5 and on 8 μm cryosections for E14.5, E16.5 and P0 (birth) gonads. The EGFP was detected in wholemount gonads by direct fluorescent imaging, and an anti-GFP antibody was used for immunofluorescences on sections. For the different knockout models, gonads were fixed in 4% paraformaldehyde overnight at 4°C. Immunofluorescence experiments were performed on 7 μm paraffin sections of E15.5 and P0 gonads as previously described^60^. The antibodies used in this study are listed in Table S1. Whole gonads and sections were imaged under a Leica DMI4000 confocal microscope.

### Real-Time PCR analysis in the mouse

For the time-course kinetics of *Runx1* expression, fetal gonads from CD-1 embryos at embryonic day E11.5, E12.5, E13.5, E14.5, E16.5, E18.5 and postnatal day P3 were separated from the mesonephros and snap-frozen. For each stage, 3 biological replicates were collected, with 6 gonads/replicate for the E11.5 stage and 3 gonads/replicate for the other stages. For *Runx1* KO analysis, control and KO ovaries were collected at E14.5 (n = 4 biological replicates/genotype). For *Runx1/Foxl2* DKO analysis, control, *Runx1* and *Foxl2* single and double KO ovaries were collected at E15.5 (n=4/genotype) and P0 (n=5/genotype). For all experiments, total RNA was isolated for each replicate using PicoPure RNA isolation kit (Arcturus, Mountain View, CA), RNA quality and concentration were determined using the Nanodrop 2000c and 300 to 400 ng of RNA was used for cDNA synthesis with the Superscript II cDNA synthesis system (Invitrogen Corp., Carlsbad, CA). Gene expression was analyzed by real-time PCR using Bio-Rad CFX96TM Real-Time PCR Detection system. Gene expression was normalized to *Gapdh*. The Taqman probes and primers used to detect transcript expression are listed in Table S1. Data were analyzed using Prism GraphPad Software by unpaired Student’s *t-test* or by ANOVA p<0.05. Values are presented as mean ± s.e.m.

### *Runx1* expression in other species

For the rainbow trout, *Runx1* expression during gonadal development was assessed by quantitative PCR, as previously described^61^. Species-specific primers used are listed in Table S1. For the red-eared slider turtle, *Runx1* expression during gonadal development was assessed at Female-Promoting Temperature (FPT) of 31°C and at Male-Promoting Temperature (MPT) of 26°C by RNA-seq, as previously described^62^. For the goat, *Runx1* expression during gonadal development was assessed by quantitative PCR, and 2 to 3 biological replicates were used for each stage of development as previously described^63^. Values are presented as mean ± s.d. All goat handling procedures were conducted in compliance with the guidelines on the Care and Use of Agricultural Animals in Agricultural Research and Teaching in France (Authorization no. 91-649 for the Principal Investigator, and national authorizations for all investigators. Approval from the Ethics Committee: 12/045). For the human, *Runx1* expression during gonadal development was assessed by RNA-seq (Lecluze *et al*. in preparation). Human fetuses (6-12 GW) were obtained from legally-induced normally-progressing terminations of pregnancy performed in Rennes University Hospital in France. Tissues were collected with women’s written consent, in accordance with the legal procedure agreed by the National agency for biomedical research (authorization #PFS09-011; Agence de la Biomédecine) and the approval of the Local ethics committee of Rennes Hospital in France (advice # 11-48).

### Microarray analysis

Gene expression analysis of control, *Runx1* KO, *Foxl2* KO and *Runx1/Foxl2* DKO ovaries was conducted using Affymetrix Mouse Genome 430 2.0 GeneChip® arrays (Affymetrix, Santa Clara, CA) on 4 biological replicates (one P0 gonad per replicate) for each genotype. Fifty (50) nanograms of total RNA were amplified and labeled as directed in the WT-Ovation Pico RNA Amplification System and Encore Biotin Module protocols. Amplified biotin-aRNA (4.6 μg) was fragmented and hybridized to each array for 18 hours at 45°C in a rotating hybridization. Array slides were stained with streptavidin/phycoerythrin utilizing a double-antibody staining procedure and then washed for antibody amplification according to the GeneChip Hybridization, Wash and Stain Kit and user manual following protocol FS450-0004. Arrays were scanned in an Affymetrix Scanner 3000 and data was obtained using the GeneChip® Command Console Software (AGCC; Version 3.2) and Expression Console (Version 1.2). Microarray data have been deposited in GEO under accession code GSE129038. Gene expression analyses were conducted with Partek software (St. Louis, Missouri) using a one-way ANOVA comparing the RMA normalized log2 intensities. A full dataset Excel file containing the normalized log2 intensity of all genes for each genotype, and a graphic view of their expression is provided in Dataset S2. In order to identify differentially expressed genes, analysis of variance (ANOVA) was used to determine if there was a statistical difference between the means of groups and the gene lists were filtered with p<0.05 and fold-change cutoff of 1.5. The heat-map was created comparing the genes that were significantly different between control and *Runx1/Foxl2* double knockout ovaries. Venn diagrams were generated in Partek by comparing gene-symbols between the lists of genes differentially expressed.

### ChIP-seq assays and analysis

Ovaries from E14.5 CD-1 embryos were separated from the mesonephros, snap-frozen, and stored at −80°C. RUNX1 ChIP-seq experiments and analyses in E14.5 ovaries were performed as previously described for FOXL2 ChIP-seq^41^. Two independent ChIP-seq experiments were performed by Active Motif Inc. using 20-30 μg of sheared chromatin from pooled embryonic ovaries (n=100-120 ovaries/ChIP), and 10 μl of RUNX1 antibody^64^ (provided by Drs. Yoram Groner and Ditsa Levanon, the Weizmann Institute of Science, Israel). ChIP-seq libraries were sequenced as single-end 75-mers by Illumina NextSeq 500, then filtered to retain only reads with average base quality score >20. Reads were mapped against the mouse mm10 reference genome using Bowtie^65^ v1.2 with parameter “-m 1” to collect only uniquely-mapped hits. Duplicate mapped reads were removed using Picard tools MarkDuplicates.jar (v1.110). The number of uniquely-mapped non-duplicate reads for each biological replicate was 8,932,674 and 15,036,698. After merging the replicate datasets, binding regions were identified by peak calling using HOMER v.4.9 ^66^ with FDR<1e-5. Called peaks were subsequently redefined as 300mers centered on the called peak midpoints and filtered for a 4-fold enrichment over input and over local signal. Genomic distribution of RUNX1-bound regions was determined based on Refseq gene models as downloaded from the UCSC Genome Browser as of August 09, 2017. Enriched motifs were identified using HOMER findMotifsGenome.pl de novo motif analysis with parameter “-size given”. For RUNX1 and FOXL2 ChIP-seq comparisons, binding peaks that had at least 1 bp in common were considered overlapping. Peaks were assigned to the nearest gene based on RefSeq. Gene lists were analyzed for enrichment using the online tool EnrichR ^67^. The ChIP-seq data are available in the ReproGenomics Viewer (https://rgv.genouest.org)^68, 69^ and Gene Expression Omnibus (GSE128767; http://www.ncbi.nlm.nih.gov/geo/).

## Acknowledgments

We thank the late Keith Parker (UT Southwestern Medical Center, USA) for the *Sf1*-Cre mice, David Schlessinger (National Institute on Aging, USA) for the *Foxl2*^+/-^ mice, Yoram Groner and Ditsa Levanon (The Weizmann Institute of Science, Israel) for the RUNX1 antibody, David Zarkower (University of Minnesota, USA) for the DMRT1 antibody, Ken Morohashi (Kyushu University, Japan) for the SF1 and SOX9 antibodies, and Dagmar Wilhelm (University of Melbourne, Australia) for the SRY antibody. We are grateful for the NIEHS Molecular Genomics Core, Integrative Bioinformatics Support Group, Comparative Medicine Branch, and Cellular and Molecular Pathology Branch for their services.

## Authors contributions

B.N. performed the experiments in the mouse; B.N. and H.H.-C.Y. designed the study, analyzed data and wrote the paper; S.A.G performed bioinformatic analyses; F.C and E.L. analyzed *RUNX1* expression in human fetal gonads; M.P. and E.P. analyzed *RUNX1* expression in the goat; E.D-D-P. and Y.G. analyzed *runx1* expression in rainbow trout; B.C. analyzed *Runx1* expression in the red-eared slider turtle. S.A.G, E.L., F.C., M.P., E.P., E.D-D-P., Y.G., B.C., and H.H-C.Y. edited the paper.

## Materials & Correspondence

Correspondence and requests for materials should be addressed to H.H-C.Y.

## References

1. Capel B. Vertebrate sex determination: evolutionary plasticity of a fundamental switch. Genetics 18, 675–689 (2017).

2. Chassot AA, et al. Activation of beta-catenin signaling by Rspo1 controls differentiation of the mammalian ovary. Hum Mol Genet 17, 1264–1277 (2008).

3. Tomizuka K, et al. R-spondin1 plays an essential role in ovarian development through positively regulating Wnt-4 signaling. Hum Mol Genet 17, 1278–1291 (2008).

4. Vainio S, Heikkila M, Kispert A, Chin N, McMahon AP. Female development in mammals is regulated by Wnt-4 signalling. Nature 397, 405–409 (1999).

5. Ottolenghi C, et al. Foxl2 is required for commitment to ovary differentiation. Hum Mol Genet 14, 2053–2062 (2005).

6. Schmidt D, et al. The murine winged-helix transcription factor Foxl2 is required for granulosa cell differentiation and ovary maintenance. Development 131, 933–942 (2004).

7. Uhlenhaut NH, et al. Somatic Sex Reprogramming of Adult Ovaries to Testes by FOXL2 Ablation. Cell 139, 1130–1142 (2009).

8. Auguste A, et al. Loss of R-spondin1 and Foxl2 amplifies female-to-male sex reversal in XX mice. Sex Dev 5, 304–317 (2011).

9. Ottolenghi C, et al. Loss of Wnt4 and Foxl2 leads to female-to-male sex reversal extending to germ cells. Hum Mol Genet 16, 2795–2804 (2007).

10. Herpin A, Schartl M. Plasticity of gene-regulatory networks controlling sex determination: of masters, slaves, usual suspects, newcomers, and usurpators. EMBO Rep 16, 1260–1274 (2015).

11. Crisponi L, et al. The putative forkhead transcription factor FOXL2 is mutated in blepharophimosis/ptosis/epicanthus inversus syndrome. Nat Genet 27, 159–166 (2001).

12. Boulanger L, et al. FOXL2 is a female sex-determining gene in the goat. Curr Biol 24, 404–408 (2014).

13. Li M, et al. Efficient and heritable gene targeting in tilapia by CRISPR/Cas9. Genetics 197, 591–599 (2014).

14. Bertho S, et al. The unusual rainbow trout sex determination gene hijacked the canonical vertebrate gonadal differentiation pathway. Proc Natl Acad Sci U S A 115, 12781–12786 (2018).

15. Raymond CS, et al. Evidence for evolutionary conservation of sex-determining genes. Nature 391, 691–695 (1998).

16. Matsuda M, et al. DMY is a Y-specific DM-domain gene required for male development in the medaka fish. Nature 417, 559–563 (2002).

17. Nanda I, et al. A duplicated copy of DMRT1 in the sex-determining region of the Y chromosome of the medaka, Oryzias latipes. Proc Natl Acad Sci U S A 99, 11778–11783 (2002).

18. Matson CK, Murphy MW, Sarver AL, Griswold MD, Bardwell VJ, Zarkower D. DMRT1 prevents female reprogramming in the postnatal mammalian testis. Nature 476, 101–104 (2011).

19. Mello MP, et al. Novel DMRT1 3’UTR+11insT mutation associated to XY partial gonadal dysgenesis. Arq Bras Endocrinol Metabol 54, 749–753 (2010).

20. Raymond CS, Murphy MW, O’Sullivan MG, Bardwell VJ, Zarkower D. Dmrt1, a gene related to worm and fly sexual regulators, is required for mammalian testis differentiation. Genes Dev 14, 2587–2595 (2000).

21. Duffy JB, Gergen JP. The Drosophila Segmentation Gene Runt Acts as a Position-Specific Numerator Element Necessary for the Uniform Expression of the Sex-Determining Gene Sex-Lethal. Genes Dev 5, 2176–2187 (1991).

22. Kramer SG, Jinks TM, Schedl P, Gergen JP. Direct activation of Sex-lethal transcription by the Drosophila Runt protein. Development 126, 191–200 (1999).

23. Nef S, et al. Gene expression during sex determination reveals a robust female genetic program at the onset of ovarian development. Dev Biol 287, 361–377 (2005).

24. Sullivan JC, et al. The evolutionary origin of the Runx/CBFbeta transcription factors--studies of the most basal metazoans. BMC Evol Biol 8, 228 (2008).

25. Groner Y, Ito Y, Liu P, Neil JC, Speck NA, van Wijnen A (eds). RUNX Proteins in Development and Cancer, Advances in Experimental Medicine and Biology 962. (Springer, 2017).

26. Chuang LS, Ito K, Ito Y. RUNX family: Regulation and diversification of roles through interacting proteins. Int J Cancer 132, 1260–1271 (2013).

27. Gong S, et al. A gene expression atlas of the central nervous system based on bacterial artificial chromosomes. Nature 425, 917–925 (2003).

28. Koopman P, Gubbay J, Vivian N, Goodfellow P, Lovell-Badge R. Male development of chromosomally female mice transgenic for Sry. Nature 351, 117–121 (1991).

29. Mork L, et al. Temporal Differences in Granulosa Cell Specification in the Ovary Reflect Distinct Follicle Fates in Mice. Biol Reprod 86, (2012).

30. Rastetter RH, et al. Marker genes identify three somatic cell types in the fetal mouse ovary. Dev Biol 394, 242–252 (2014).

31. Okuda T, vanDeursen J, Hiebert SW, Grosveld G, Downing JR. AML1, the target of multiple chromosomal translocations in human leukemia, is essential for normal fetal liver hematopoiesis. Cell 84, 321–330 (1996).

32. Wang Q, Stacy T, Binder M, MarinPadilla M, Sharpe AH, Speck NA. Disruption of the Cbfa2 gene causes necrosis and hemorrhaging in the central nervous system and blocks definitive hematopoiesis. P Natl Acad Sci USA 93, 3444–3449 (1996).

33. Bingham NC, Verma-Kurvari S, Parada LF, Parker KL. Development of a steroidogenic factor 1/Cre transgenic mouse line. Genesis 44, 419–424 (2006).

34. Lei N, Hornbaker KI, Rice DA, Karpova T, Agbor VA, Heckert LL. Sex-specific differences in mouse DMRT1 expression are both cell type- and stage-dependent during gonad development. Biol Reprod 77, 466–475 (2007).

35. Chaboissier MC, et al. Functional analysis of Sox8 and Sox9 during sex determination in the mouse. Development 131, 1891–1901 (2004).

36. Vidal VP, Chaboissier MC, de Rooij DG, Schedl A. Sox9 induces testis development in XX transgenic mice. Nat Genet 28, 216–217 (2001).

37. Jorgez CJ, Klysik M, Jamin SP, Behringer RR, Matzuk MM. Granulosa cell-specific inactivation of follistatin causes female fertility defects. Mol Endocrinol 18, 953–967 (2004).

38. Britt KL, et al. The ovarian phenotype of the aromatase knockout (ArKO) mouse. J Steroid Biochem Mol Biol 79, 181–185 (2001).

39. Jameson SA, et al. Temporal transcriptional profiling of somatic and germ cells reveals biased lineage priming of sexual fate in the fetal mouse gonad. PLoS Genet 8, e1002575 (2012).

40. Colvin JS, Green RP, Schmahl J, Capel B, Ornitz DM. Male-to-female sex reversal in mice lacking fibroblast growth factor 9. Cell 104, 875–889 (2001).

41. Nicol B, Grimm SA, Gruzdev A, Scott GJ, Ray MK, Yao HHC. Genome-wide identification of FOXL2 binding and characterization of FOXL2 feminizing action in the fetal gonads. Hum Mol Genet 27, 4273–4287 (2018).

42. Ong CT, Corces VG. CTCF: an architectural protein bridging genome topology and function. Nature Reviews Genetics 15, 234–246 (2014).

43. Park M, et al. FOXL2 interacts with steroidogenic factor-1 (SF-1) and represses SF-1- induced CYP17 transcription in granulosa cells. Mol Endocrinol 24, 1024–1036 (2010).

44. Yang WH, Gutierrez NM, Wang LZ, Ellsworth BS, Wang CM. Synergistic Activation of the Mc2r Promoter by FOXL2 and NR5A1 in Mice. Biol Reprod 83, 842–851 (2010).

45. Meyers S, Downing JR, Hiebert SW. Identification of AML-1 and the (8;21) translocation protein (AML-1/ETO) as sequence-specific DNA-binding proteins: the runt homology domain is required for DNA binding and protein-protein interactions. Mol Cell Biol 13, 6336–6345 (1993).

46. Blount AL, Schmidt K, Justice NJ, Vale WW, Fischer WH, Bilezikjian LM. FoxL2 and Smad3 coordinately regulate follistatin gene transcription. J Biol Chem 284, 7631–7645 (2009).

47. Barrionuevo F, et al. Homozygous inactivation of Sox9 causes complete XY sex reversal in mice. Biol Reprod 74, 195–201 (2006).

48. Wildey GM, Howe PH. Runx1 is a co-activator with FOXO3 to mediate transforming growth factor beta (TGFbeta)-induced Bim transcription in hepatic cells. J Biol Chem 284, 20227–20239 (2009).

49. Wang L, Brugge JS, Janes KA. Intersection of FOXO- and RUNX1-mediated gene expression programs in single breast epithelial cells during morphogenesis and tumor progression. Proc Natl Acad Sci U S A 108, E803–812 (2011).

50. Ono M, et al. Foxp3 controls regulatory T-cell function by interacting with AML1/Runx1. Nature 446, 685–689 (2007).

51. Recouvreux MS, et al. RUNX1 and FOXP3 interplay regulates expression of breast cancer related genes. Oncotarget 7, 6552–6565 (2016).

52. Munger SC, Natarajan A, Looger LL, Ohler U, Capel B. Fine time course expression analysis identifies cascades of activation and repression and maps a putative regulator of mammalian sex determination. PLoS Genet 9, e1003630 (2013).

53. Stevant I, et al. Deciphering Cell Lineage Specification during Male Sex Determination with Single-Cell RNA Sequencing. Cell reports 22, 1589–1599 (2018).

54. Nottingham WT, et al. Runx1-mediated hematopoietic stem-cell emergence is controlled by a Gata/Ets/SCL-regulated enhancer. Blood 110, 4188–4197 (2007).

55. Albrecht KH, Eicher EM. Evidence that Sry is expressed in pre-Sertoli cells and Sertoli and granulosa cells have a common precursor. Dev Biol 240, 92–107 (2001).

56. Feng L, et al. ADAM10-Notch signaling governs the recruitment of ovarian pregranulosa cells and controls folliculogenesis in mice. J Cell Sci 129, 2202–2212 (2016).

57. Rahmoun M, et al. In mammalian foetal testes, SOX9 regulates expression of its target genes by binding to genomic regions with conserved signatures. Nucleic Acids Res 45, 7191–7211 (2017).

58. Levasseur A, Paquet M, Boerboom D, Boyer A. Yes-associated protein and WW-containing transcription regulator 1 regulate the expression of sex-determining genes in Sertoli cells, but their inactivation does not cause sex reversal. Biol Reprod 97, 162–175 (2017).

59. Uda M, et al. Foxl2 disruption causes mouse ovarian failure by pervasive blockage of follicle development. Hum Mol Genet 13, 1171–1181 (2004).

60. Nicol B, Yao HH. Gonadal Identity in the Absence of Pro-Testis Factor SOX9 and Pro-Ovary Factor Beta-Catenin in Mice. Biol Reprod 93, 35 (2015).

61. Marivin E, et al. Sex hormone-binding globulins characterization and gonadal gene expression during sex differentiation in the rainbow trout, Oncorhynchus mykiss. Mol Reprod Dev 81, 757–765 (2014).

62. Czerwinski M, Natarajan A, Barske L, Looger LL, Capel B. A timecourse analysis of systemic and gonadal effects of temperature on sexual development of the red-eared slider turtle Trachemys scripta elegans. Dev Biol 420, 166–177 (2016).

63. Elzaiat M, et al. High-throughput sequencing analyses of XX genital ridges lacking FOXL2 reveal DMRT1 up-regulation before SOX9 expression during the sex-reversal process in goats. Biol Reprod 91, 153 (2014).

64. Umansky KB, et al. Runx1 Transcription Factor Is Required for Myoblasts Proliferation during Muscle Regeneration. PLoS Genet 11, e1005457 (2015).

65. Langmead B, Trapnell C, Pop M, Salzberg SL. Ultrafast and memory-efficient alignment of short DNA sequences to the human genome. Genome Biol 10, R25 (2009).

66. Heinz S, et al. Simple combinations of lineage-determining transcription factors prime cis-regulatory elements required for macrophage and B cell identities. Mol Cell 38, 576–589 (2010).

67. Chen EY, et al. Enrichr: interactive and collaborative HTML5 gene list enrichment analysis tool. BMC Bioinformatics 14, 128 (2013).

68. Darde TA, et al. The ReproGenomics Viewer: a multi-omics and cross-species resource compatible with single-cell studies for the reproductive science community. Bioinformatics, (2019).

69. Darde TA, et al. The ReproGenomics Viewer: an integrative cross-species toolbox for the reproductive science community. Nucleic Acids Res 43, W109–116 (2015).

