## Supplementary Figures for "RUNX1 safeguards the identity of the fetal ovary through an interplay with FOXL2"

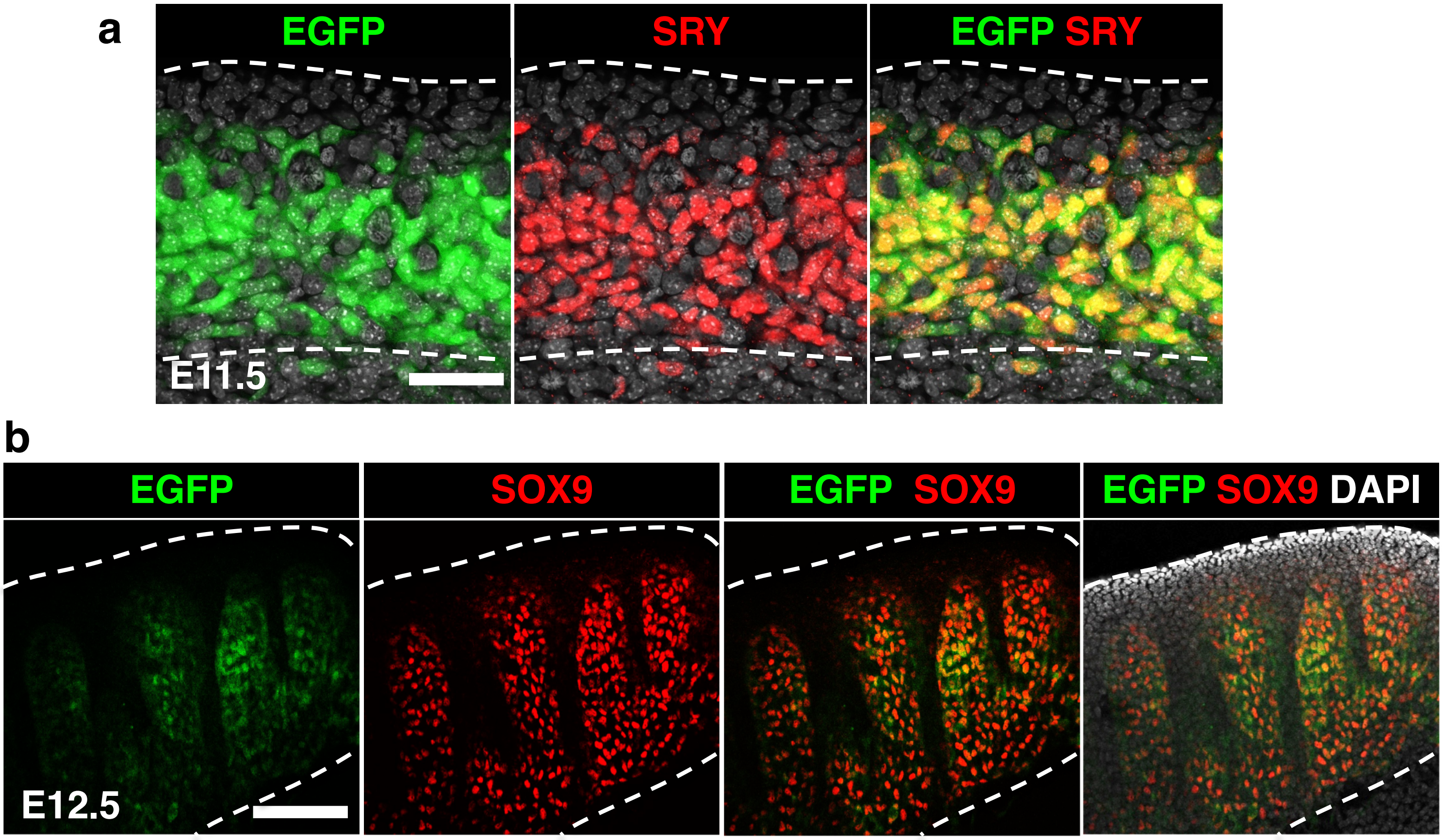


**Figure S1: *Runx1* expression in the fetal testis. (a)** Whole-mount immunofluorescence for SRY in XY Tg(Runx1-EGFP) gonads at E11.5. Scale bar: 50 $\mu$m. *Runx1*-EGFP is exclusively expressed in SRY+ supporting cells. **(b)** Whole-mount immunofluorescence for SOX9 in XY Tg(Runx1-EGFP) gonads at E12.5. Scale bar: 100 $\mu$m. Low levels of EGFP are detected in SOX9+ cells.

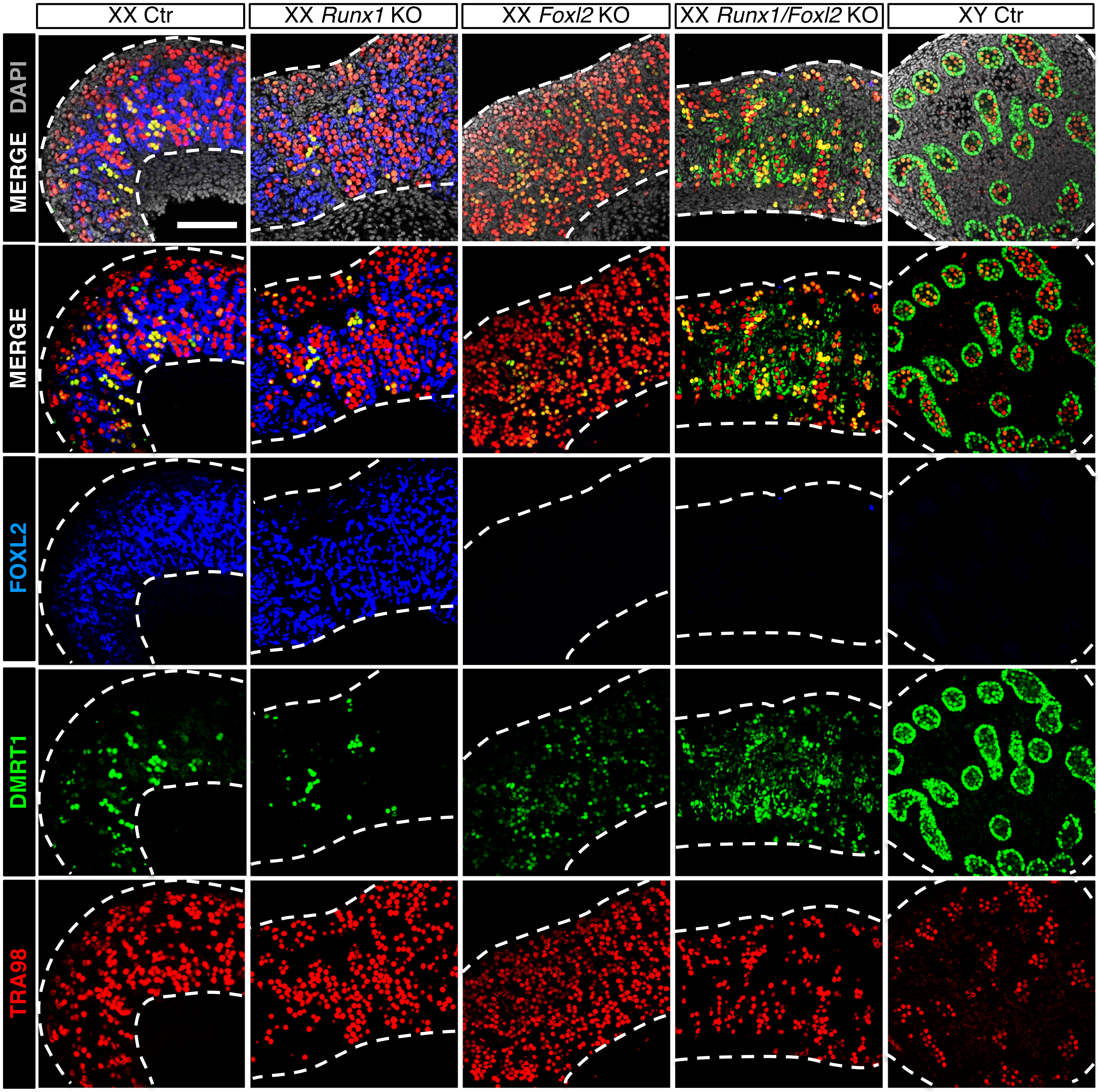


**Figure S2: Single channels for the FOXL2/DMRT1/TRA98 immunofluorescences at E15.5 shown in Fig. 5 a-e.** Scale bar: 100 $\mu$m. Dotted lines outline the gonads.

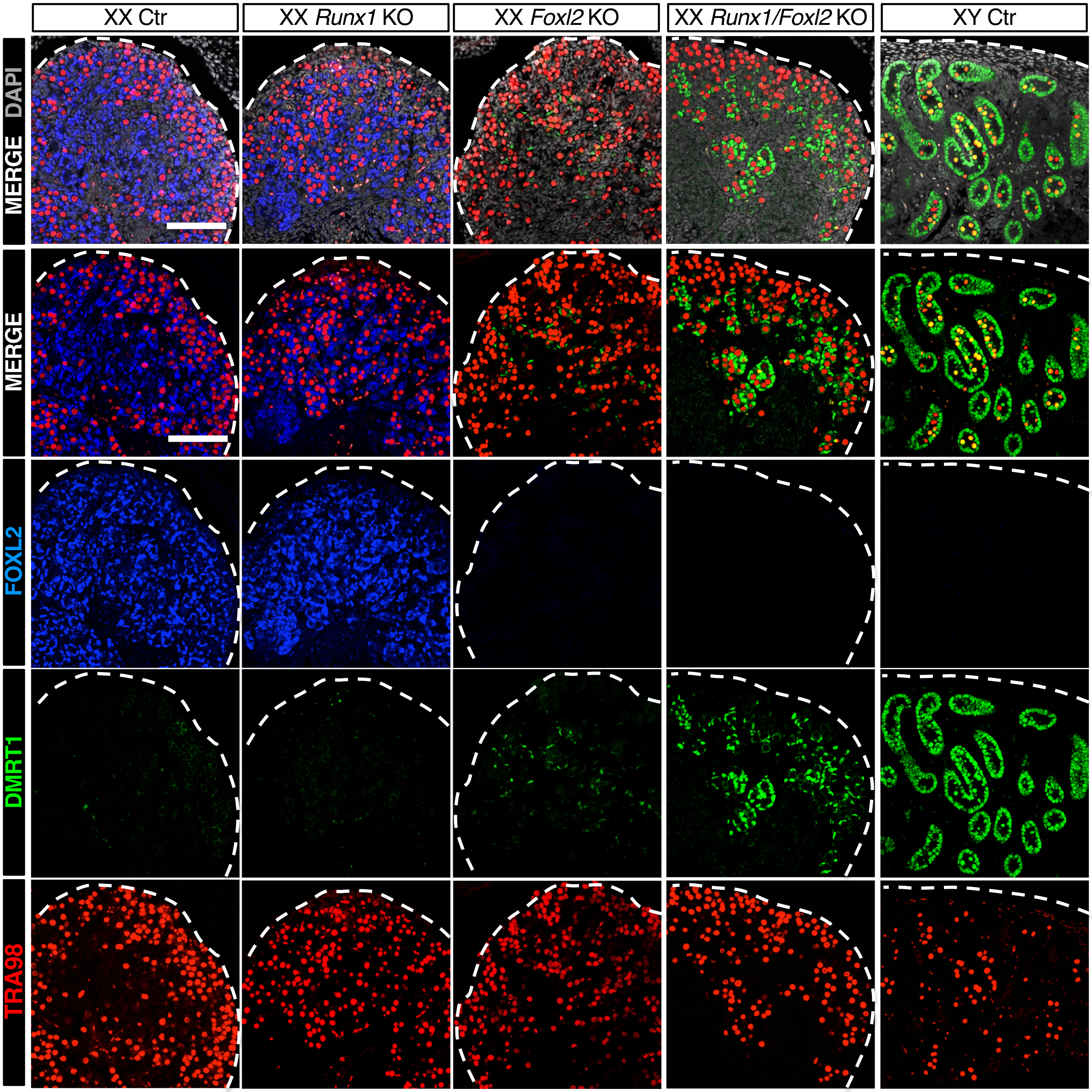


**Figure S3: Single channels for the FOXL2/DMRT1/TRA98 immunofluorescence at birth shown in Fig. 5 f-j.** Scale bar: 100 $\mu$m. Dotted lines outline the gonads.

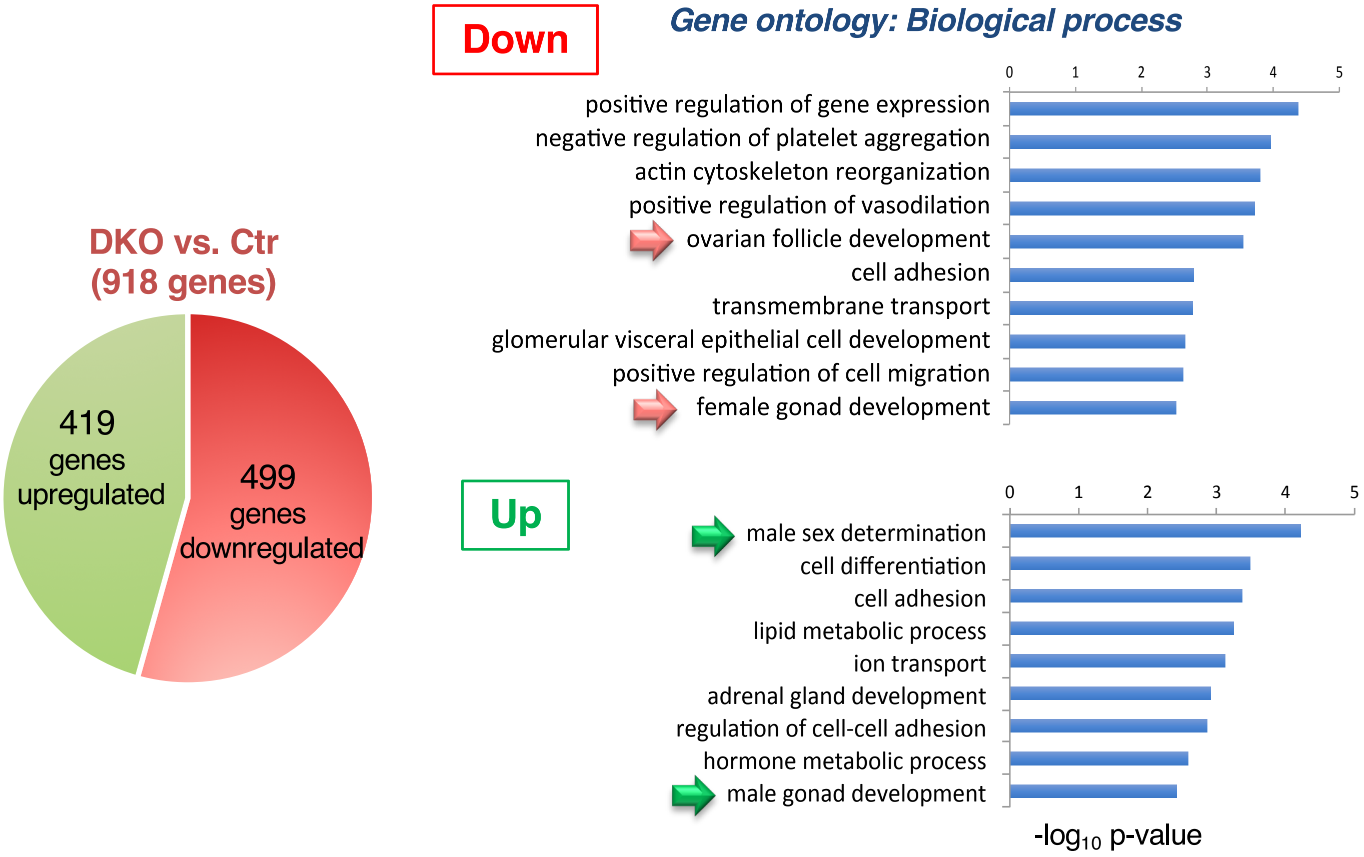


**Figure S4: Gene ontology biological process analysis of genes downregulated or upregulated in *Runx1/Foxl2* DKO vs. control ovaries at birth.** Top GO-terms are shown. Analysis performed using DAVID 6.8 on genes significantly changed with a fold-change >1.5 and *P*<0.05.
